# Visualizing Population Structure with Variational Autoencoders

**DOI:** 10.1101/2020.08.12.248278

**Authors:** C. J. Battey, Gabrielle C. Coffing, Andrew D. Kern

## Abstract

Dimensionality reduction is a common tool for visualization and inference of population structure from genotypes, but popular methods either return too many dimensions for easy plotting (PCA) or fail to preserve global geometry (t-SNE and UMAP). Here we explore the utility of variational autoencoders (VAEs) – generative machine learning models in which a pair of neural networks seek to first compress and then recreate the input data – for visualizing population genetic variation. VAEs incorporate non-linear relationships, allow users to define the dimensionality of the latent space, and in our tests preserve global geometry better than t-SNE and UMAP. Our implementation, which we call `popvae`, is available as a command-line python program at github.com/kr-colab/popvae. The approach yields latent embeddings that capture subtle aspects of population structure in humans and *Anopheles* mosquitoes, and can generate artificial genotypes characteristic of a given sample or population.

## Introduction

As we trace the genealogy of a population forward in time, branching inherent in the genealogical process leads to hierarchical relationships among individuals that can be thought of as clades. Much of the genetic variation among individuals in a species thus reflects the history of isolation and migration of their ancestors. Describing this population structure is itself a major goal in biogeography, systematics, and human genetics; wherein one might attempt to infer the number of genotypic clusters supported by the data (Holsinger and Weir, 2009), estimate relative rates of migration (Petkova et al., 2016), or observe turnover in the ancestry of people living in a geographic region (Antonio et al., 2019).

Estimation of population structure is also critical for our ability to accurately link genetic variation to phenotypic variation, because population structure is a major confounding factor in genome-wide association studies (GWAS) (Lander and Schork, 1994; Pritchard and Donnelly, 2001; Marchini et al., 2004; Freedman et al., 2004). Downstream studies that use GWAS information can themselves be compromised by inadequate controls for structure, for instance in recent work trying to identify the effects of natural selection on complex traits (Mathieson and McVean, 2012; Berg et al., 2019; Sohail et al., 2019). Dimensionality reduction via principal components analysis (PCA) has been an important tool for geneticists in this regard, and is now commonly used both to control for the effects of population structure in GWAS(Price et al., 2006; Patterson et al., 2006) as well as for visualization of genetic variation.

As a visualization tool however, PCA scatterplots can be difficult to interpret because information about genetic variation is split across many axes, while efficient plotting is restricted to two dimensions. Though techniques like plotting marginal distributions as stacked density plots can aid interpretation, these require binning samples into “populations” prior to visualization, are rarely used in practice, and remain difficult to interpret in complex cases. Recently two techniques from the machine learning community – t-SNE (Maaten and Hinton, 2008) and UMAP (McInnes et al., 2018) – have shown promising performance in producing two-dimensional visualizations of high-dimensional biological data. In the case of UMAP, Diaz-Papkovich et al. (2019) recently showed that running the algorithm on a large set of principal component axes allows visualization of subtle aspects of population structure in three human genotyping datasets.

However, interpreting UMAP and t-SNE plots is also complicated by a lack of so-called global structure. Though these methods perform well in clustering similar samples, distances between groups are not always meaningful – two clusters separated by a large distance in a t-SNE plot can be more similar to each other than either is to their immediate neighbors (Becht et al., 2019). The degree to which initialization and hyperparameter tuning can alleviate this issue remains an open question in the literature (Kobak and Linderman, 2019).

To create meaningful and interpretable visualizations of population genetic data we would like a method that encodes as much information as possible into just two dimensions while maintaining global structure. One way of achieving this is with a variational autoencoder (VAE).

VAEs consist of a pair of deep neural networks in which the first network (the encoder) encodes input data as a probability distribution in a latent space and the second (the decoder) seeks to recreate the input given a set of latent coordinates (Kingma and Welling, 2013). Thus a VAE has as its target the input data itself. The loss function for a VAE is the sum of reconstruction error (how different the generated data is from the input) and Kullback-Leibler (KL) divergence between a sample’s distribution in latent space and a reference distribution which acts as a prior on the latent space (here we use a standard multivariate normal, but see (Davidson et al., 2018) for an alternative design with a hyperspherical latent space). The KL term of the loss function incentivizes the encoder to generate latent distributions with meaningful distances among samples, while the reconstruction error term helps to achieve good local clustering and data generation. VAE’s have been used extensively in image generation (Gulrajani et al., 2016; Larsen et al., 2015; Hou et al., 2016) and several recent studies have applied them to dimensionality reduction and classification of single-cell RNAseq data (Wang and Gu, 2018; Grønbech et al., 2018; Lafarge et al., 2018; Hu and Greene, 2019). At deeper timescales than we test here, Derkarabetian et al. (2019) recently explored the use of VAEs in species delimitation.

In population genetics two recent studies have studied the utility of generative deep neural networks for creating simulated genotypes. Montserrat et al. (2019) use a class-conditional VAE to generate artificial human genotypes, while Yelmen et al. (2019) use a restricted Boltzman machine and provide an in-depth assessment of the population genetic characteristics of their artificial genotypes. These studies found that such generative methods can produce short stretches of artificial genotypes that are difficult to distinguish from real data, but performance was improved by using a generative adversarial network (GAN) – either in combination with a VAE as in Montserrat et al. (2019) or as a standalone method in Yelmen et al. (2019). In this study we focus not on generation of simulated genotypes, but instead on the learned latent space representations of genotypes produced by a VAE, and study when and how they can best be used for visualizing population structure.

We introduce a new method, `popvae` (for population VAE), a command-line python program that takes as input a set of unphased genotypes and outputs sample coordinates in a low-dimensional latent space. We test `popvae` with simulated data and demonstrate its utility in empirical datasets of humans and *Anopheles* mosquitoes. In general `popvae` is most useful for complex samples for which PCA projects important aspects of structure across many axes. Relative to t-SNE and UMAP, the approach appears to better preserve global geometry at the cost of less pronounced clustering of individual sample localities. However, we show that hyperparameter tuning and stochasticity associated with train/test splits and parameter initialization are ongoing challenges for a VAE-based method, and the approach is much more computationally intensive than PCA.

## Methods

### Model

In this manuscript we describe the application of a Variational Auto-Encoder (VAE) to population genetic data for clustering and visualization Kingma and Welling (2013). Formally let *X* be our dataset consisting of *N* observations (i.e. individual genotypes) such that *X* = {*x*_1_, *x*_2_, …, *x*_𝒩_}, and let the probability of those data with some set of parameters *θ* be *p*_*θ*_(*X*). For VAEs we are interested in representing the data with a latent model, assigning some latent process parameters *z*, such that we can write a generative latent process as *p*_*θ*_(*x, z*) = *p*_*θ*_(*z*)*p*_*θ*_(*x*|*z*), where *p*_*θ*_(*z*) is the prior distribution on *z*. The last conditional probability here *p*_*θ*_(*x*|*z*) is often referred to as the decoder, as it maps from latent space to data space.

For VAEs we also define a so-called encoder model *q*_*ϕ*_(*z*|*x*), where *ϕ* represents the parameters of the encoding (the mapping of *x* to the latent space *z*), and we seek to optimize the encoder such that *q*_*ϕ*_(*z*|*x*) ≈ *p*_*θ*_(*z*|*x*). In practice the parameters *ϕ* represent the weights and biases of the encoding neural network. We thus step from data space by using

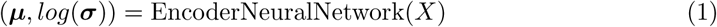

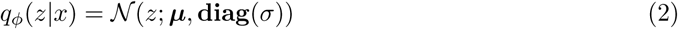

The complete VAE information flow then has three steps: the encoder estimates sample distributions in latent space as *q*_*ϕ*_(*z*|*x*), we sample from the prior on the latent space using *p*_*θ*_(*z*), and finally decode back to data space using *p*_*θ*_(*x*|*z*). Training is then performed by optimizing the *evidence lower bound* or ELBO which has parameters of the encoder and decoder within it such that

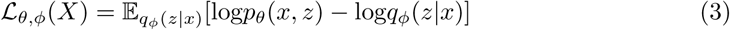

Optimization of the ELBO here leads to simultaneous fitting of the parameters of the encoder, *ϕ*, and the decoder, *θ*. In practice we use binary cross-entropy between true and generated sequences for the first term, and Kullback-Leibler divergence of sample latent distributions (relative to a standard normal 𝒩(0, 1)) for the second term of equation 3. A graphical depiction of this computational flow can be seen in Figure 1.

**Figure 1:**
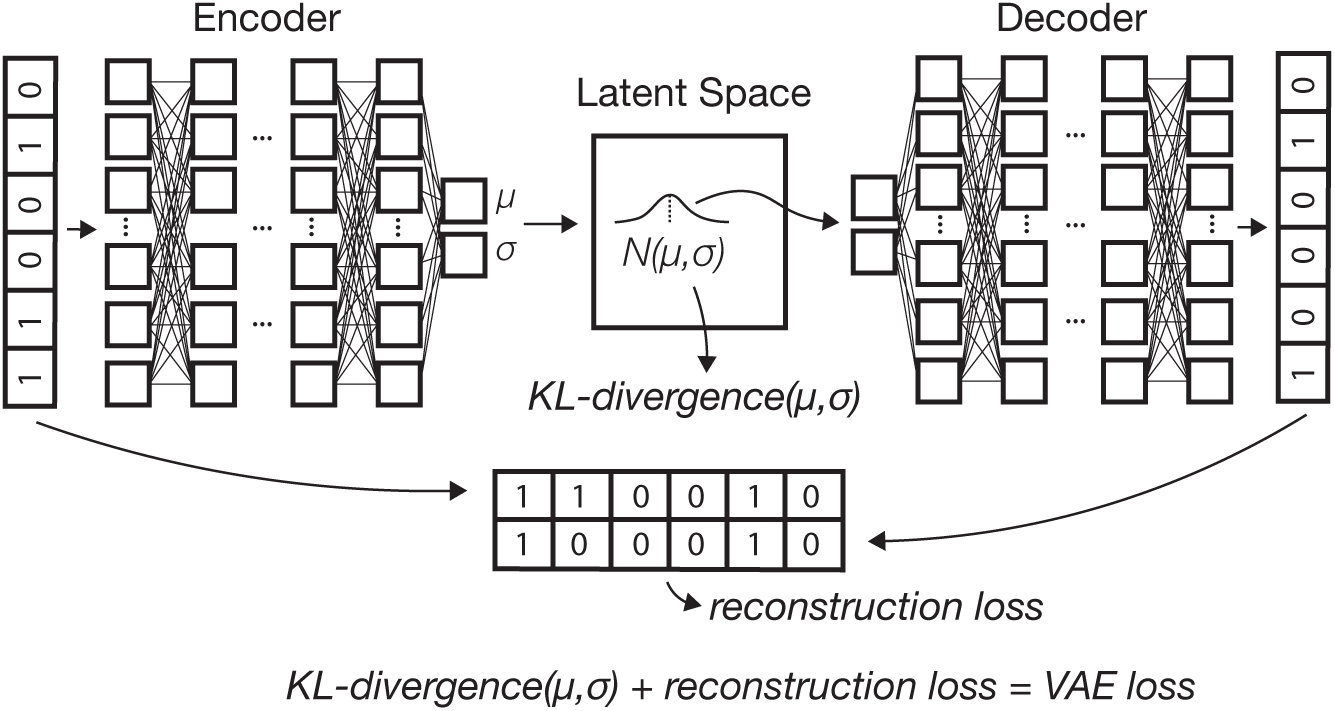
A schematic of the variational autoencoder (VAE) architecture. Input allele counts are passed to an encoder network which outputs parameters describing a sample’s location as a multivariate normal in latent space. Samples from this distribution are then passed to a decoder network which generates a new genotype vector. The loss function used to update weights and biases of both networks is the sum of reconstruction error (from comparing true and generated genotypes) and Kullback-Leibler divergence between sample latent distributions and 𝒩(0, 1).

### Implementation

We implemented this model in python 3 using the tensorflow and keras libraries (Abadi et al., 2015; Chollet et al., 2015), with preprocessing relying on numpy, pandas, and scikit-allel (Miles and Harding, 2017; Oliphant, 2006–; McKinney, 2010). `popvae` reads in genotypes from VCFs, Zarr files https://zarr.readthedocs.io/en/stable/, or a bespoke hdf5 file format. Genotypes are first filtered to remove singletons and non-biallelic sites, and missing data is filled by taking two draws from a binomial distribution with probability equal to the allele frequency across all samples (a binned version of the common practice of filling missing genotypes with the mean allele frequency (Jombart, 2008; Dray and Josse, 2015)). Filtered genotpes are then encoded with 0/0.5/1 representing homozygous ancestral, heterozygous, and homozygous derived states, respectively.

Samples are split into training and validation sets before model training. We also experimented with using all samples for training and a fixed number of epochs but found this generally led to poor performance (Appendix 1, Figure S1). Training samples are used to optimize weights and biases of the neural network, while validation samples are used to measure validation loss after each training epoch (a complete pass through the data), which in turn tunes hyperparameters of the optimizer. By default we use a random 90% of samples for training. However we found considerable variation in latent representations of some datasets when using different sets of training and validation samples (see e.g. Figure S2), so we encourage users to compare multiple training runs with different starting seeds when interpreting plots.

`popvae`’s encoder and decoder networks are fully-connected feed-forward networks whose size is controlled by two parameters – ‘`width`’, which sets the number of hidden units per layer, and ‘`depth`’, which sets the number of hidden layers. We experimented with a range of network sizes and set defaults to depth 6 and width 128, which performed well on the empirical analyses described here (Table S1, Figure S3). However we also include a grid search function by which `popvae` will conduct short training runs across a user-defined range of network sizes and then fit a final model using the network size with minimum validation loss.

We use a linear activation on the input layers to both networks and a sigmoid activation on the output of the decoder (this produces numeric values bound by (0, 1)). We interpret the sigmoid decoder outputs as the probability of observing a derived allele at a site, consistent with our 0/0.5/1 encoding of the input genotypes. All other layers use “elu” activations (Clevert et al., 2015), a modification of the more common “relu” activation which avoids the “stuck neuron” problem by returning small but nonzero values with negative inputs.

We use the Adam optimizer (Kingma and Ba, 2014) and continue model training until validation loss has not improved for *p* epochs, where *p* is a user-adjustable ‘`patience`’ parameter. We also set a learning rate scheduler to decrease the learning rate of the optimizer by half when validation loss has not improved for *p/*4 epochs. This is intended to force the optimizer to take small steps when close to the final solution, which increases training time but in our experience leads to better fit models. Users can adjust many hyperparameters from the command line, and modifying our network architectures is straightforward for those familiar with the Keras library.

To evaluate model training `popvae` returns plots of training and validation loss by epoch (e.g., Figure S4), and also outputs estimated latent coordinates for validation samples given the encoder parameters at the end of each epoch. These can then be plotted to observe how the model changes over the course of training, which can sometimes help to diagnose overfitting. We also include an interactive plotting function which generates a scatter plot of the latent space and allows users to mouse-over points to view metadata (Figure S5). This is intended to allow users to quickly iterate through models while adjusting hyperparameters. In Appendix 1 we discuss alternate approaches to network design and optimization tested while developing `popvae`.

`popvae` is available at https://github.com/kr-colab/popvae, and scripts for re-producing plots and analyses in this manuscript are available at https://github.com/cjbattey/popvae_analysis_scripts. HGDP genotypes used in this paper are available at ftp://ngs.sanger.ac.uk/production/hgdp, AG1000G genotypes at https://www.malariagen.net/data/ag1000g-phase-2-ar1, and 1000 genomes phase 3 data at https://www.internationalgenome.org/category/phase-3/.

## Results

### Latent Spaces Reflect Human Migration History

We first applied `popvae` to 100,000 SNPs from chromosome 1 in the Human Genetic Diversity Project (HGDP; Bergström et al. (2019)), a sample of global modern human diversity. The resulting latent space reflects geography from the point of view of human demographic history (Figure 2, Figure S6, Figure 4). Sub-Saharan African and South American populations are placed on opposite ends of one latent dimension, and north African (Mozabite) and east Asian samples are on opposite ends of the second; mirroring the geography of Africa and Eurasia. Samples from the Americas are roughly centered among Eurasian samples on latent dimension (LD) 2, consistent with recent demographic modeling studies suggesting a mix of Eurasian ancestries in ancestral American populations (Flegontov et al., 2019; Posth et al., 2018). Indeed the closest American samples to the European cluster are Maya individuals who were found to have low levels of recent European admixture in previous analyses(Bergström et al., 2019; Rosenberg et al., 2002) (Figure S6), suggesting `popvae` is picking up on the signal of gene flow associated with European colonization of the Americas.

**Figure 2:**
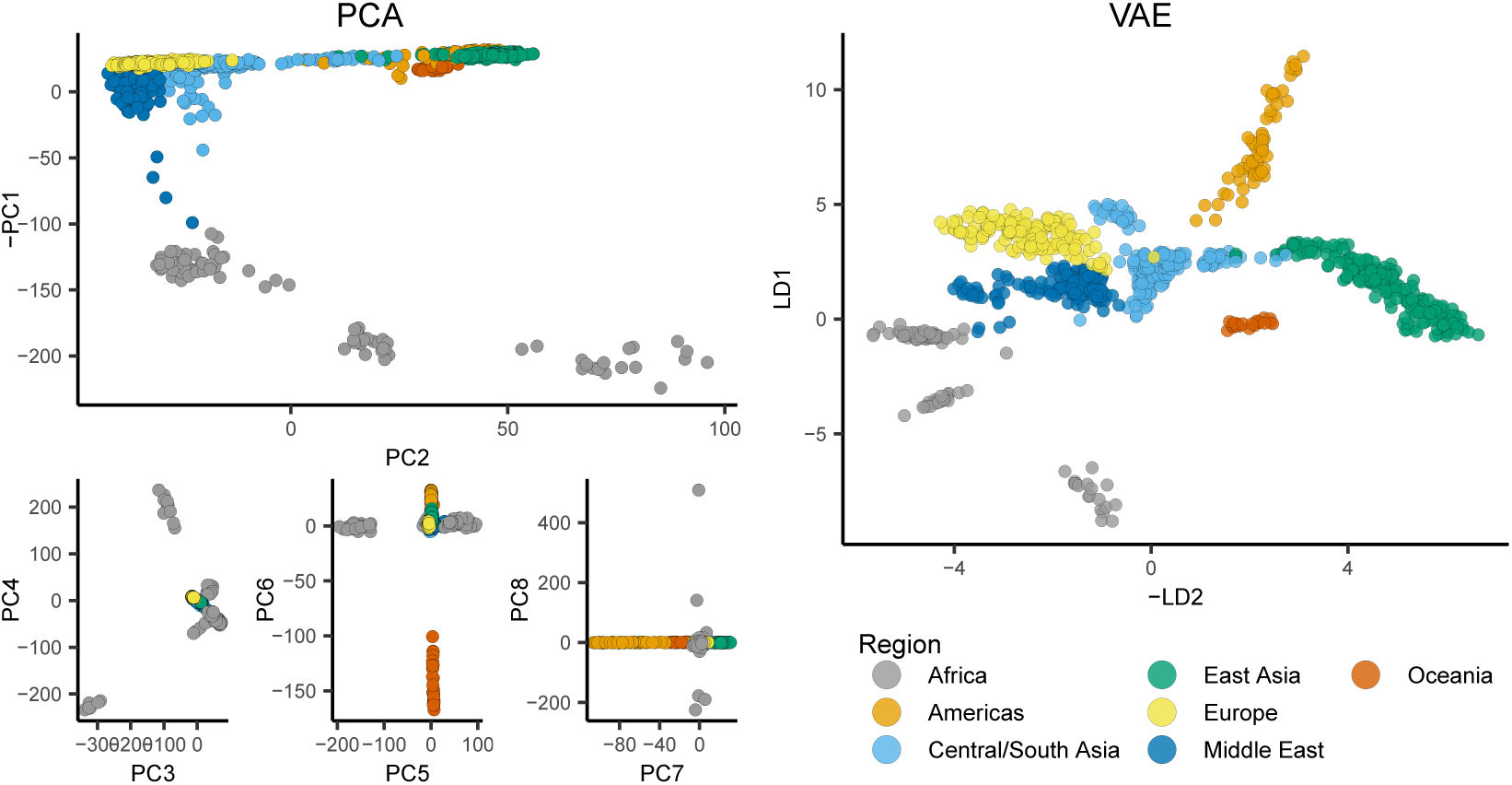
PCA axes 1-8 (left) and `popvae` run at default settings (right) for 100,000 random SNPs from chromosome 1 of the HGDP data. Axes are flipped to approximate geography.

**Figure 3:**
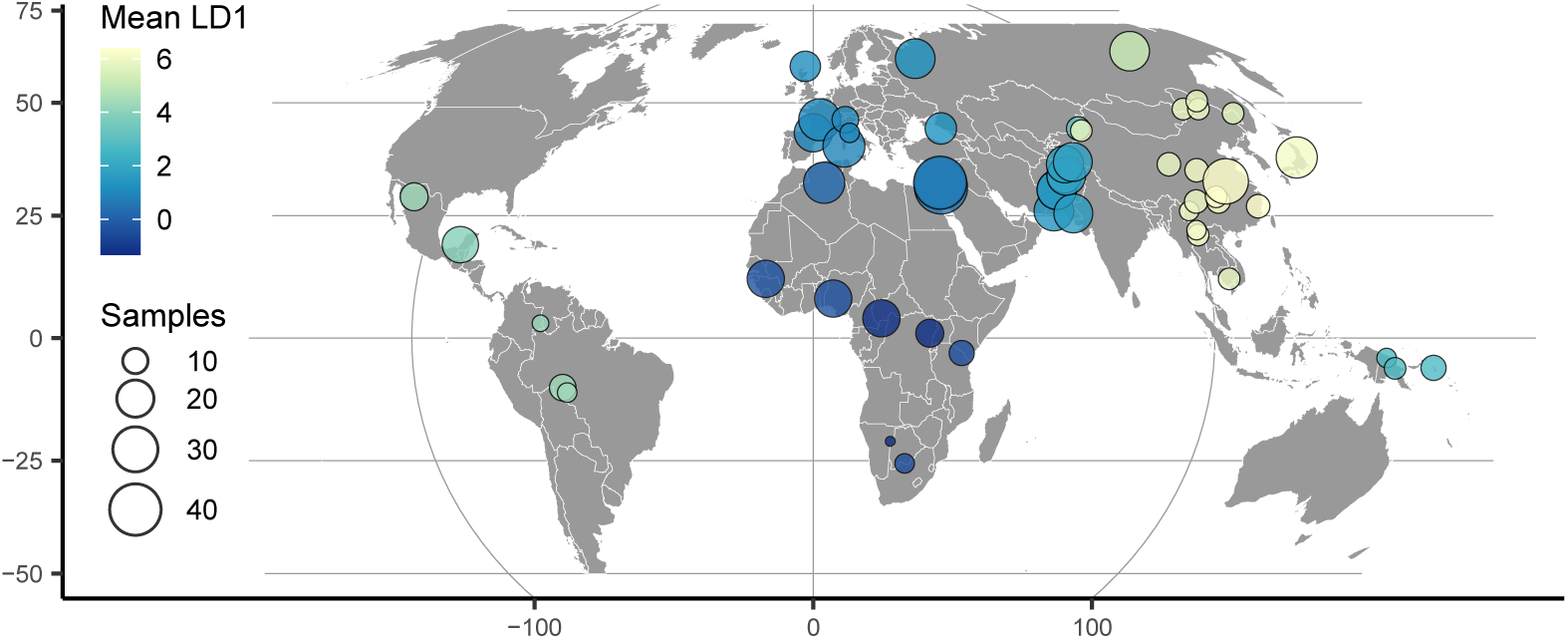
HGDP population locations with color scaled to the mean latent coordinate of a 1-dimensional `popvae` latent space.

**Figure 4:**
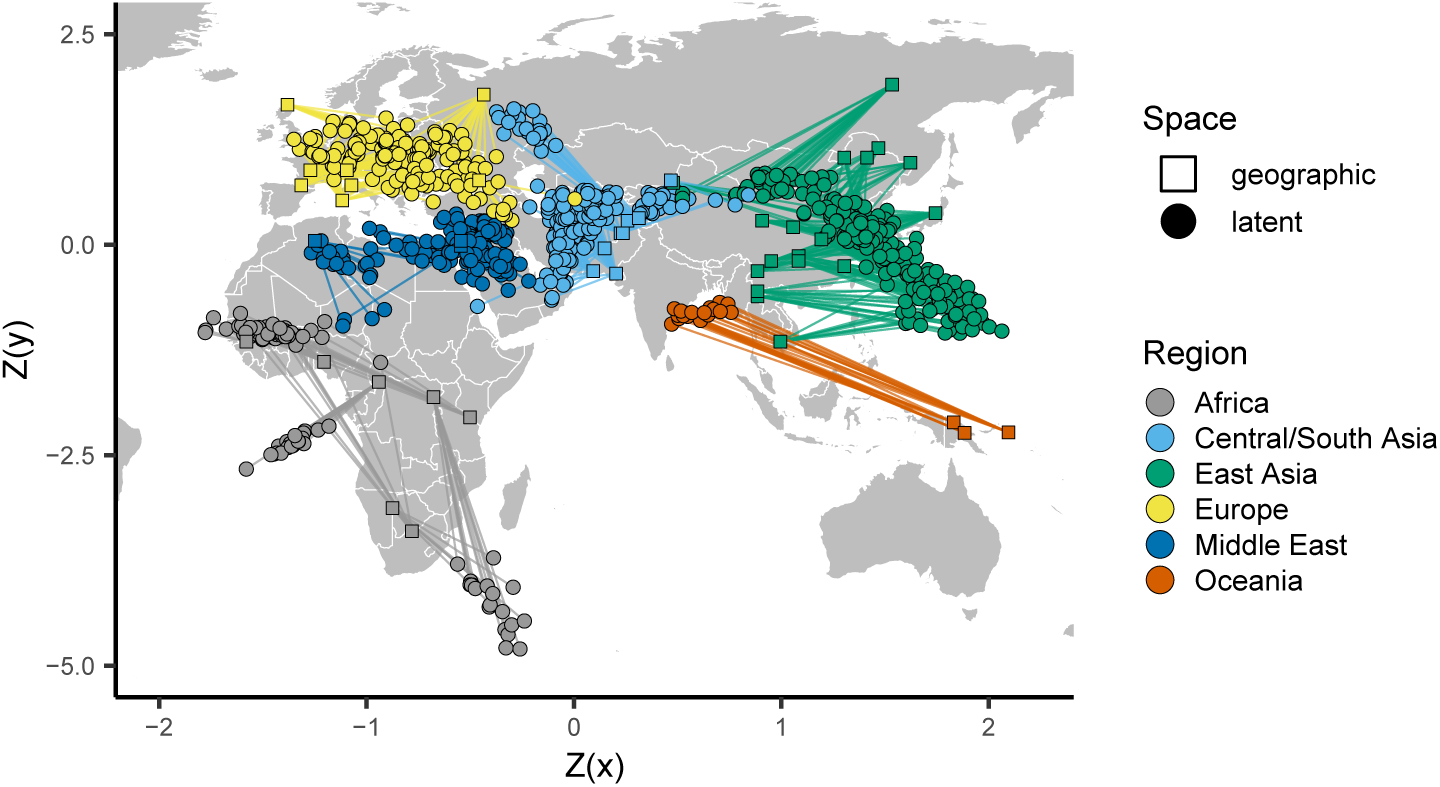
Comparing the VAE latent space with the geography of sampling localities in non-American HGDP samples (see Figure S8 for a plot including the Americas). Circles show z-normalized sample locations in latent space and squares show the corresponding location in geographic space.

These patterns are similar to those seen in PCA, but many aspects of ancestry that are difficult to see on the first two PC axes are conveniently summarized in `popvae`’s latent space. For example, differentiation within the Americas and Oceania is not visible until PC6 and PC7, respectively, but is clear in the 2D VAE latent space. This shows adjacent clusters for the islands of Bougainville and Papua New Guinea, and a cline in Eurasian ancestry from North through South America (Figure S6).

To highlight the flexibility of the VAE approach, we also trained a model with a 1-dimensional latent space and used this to scale colors on a sampling map (Figure 3). This results in a single latent dimension that approximates the diagonal of our 2D model, with African and East Asian samples on either end of the spectrum. A comparison using PCA but summarizing only the first principal component emphasizes diversity within Africa (Figure S7) and provides little resolution for out-of-Africa groups.

Finally, to emphasize the correspondence of the VAE latent space with geography, we can also directly compare geographic and latent spaces by rescaling both sets of coordinates with a z-normalization and plotting them together on a map (Figure 4). As can be seen, the visual correspondence between geographic and latent coordinates is striking in this case.

### Inversions and Population Structure in *Anopheles* Mosquitoes

We next applied `popvae` to DNA sequenced from the *Anopheles gambiae / coluzzii* complex across sub-saharan African by the AG1000G project (AG1000G Consortium, 2020; Miles et al., 2017) (Figure 5). Using 100,000 randomly-selected SNPs from chromosome 3R we again find that the VAE captures elements of population structure that are not apparent by visualizing two PC axes at a time. For example, samples from Kenya and the island of Mayotte off East Africa are highly differentiated (*F*_*st*_ > 0.18 relative to all other groups), but are placed between clusters of primarily west-African *coluzzii* and *gambiae* samples on a plot of PC1/2. The VAE instead places these populations on the opposite end of one latent dimension from all other groups and closest to Ugandan samples – similar to their relative geographic position and positions on PC3/4. The VAE also captures the relatively high differentiation of samples from Gabon and significant variation within Cameroon, which are not visible until PC6 and PC8, respectively. Further details of population structure in this species complex are discussed in AG1000G Consortium (2020).

**Figure 5:**
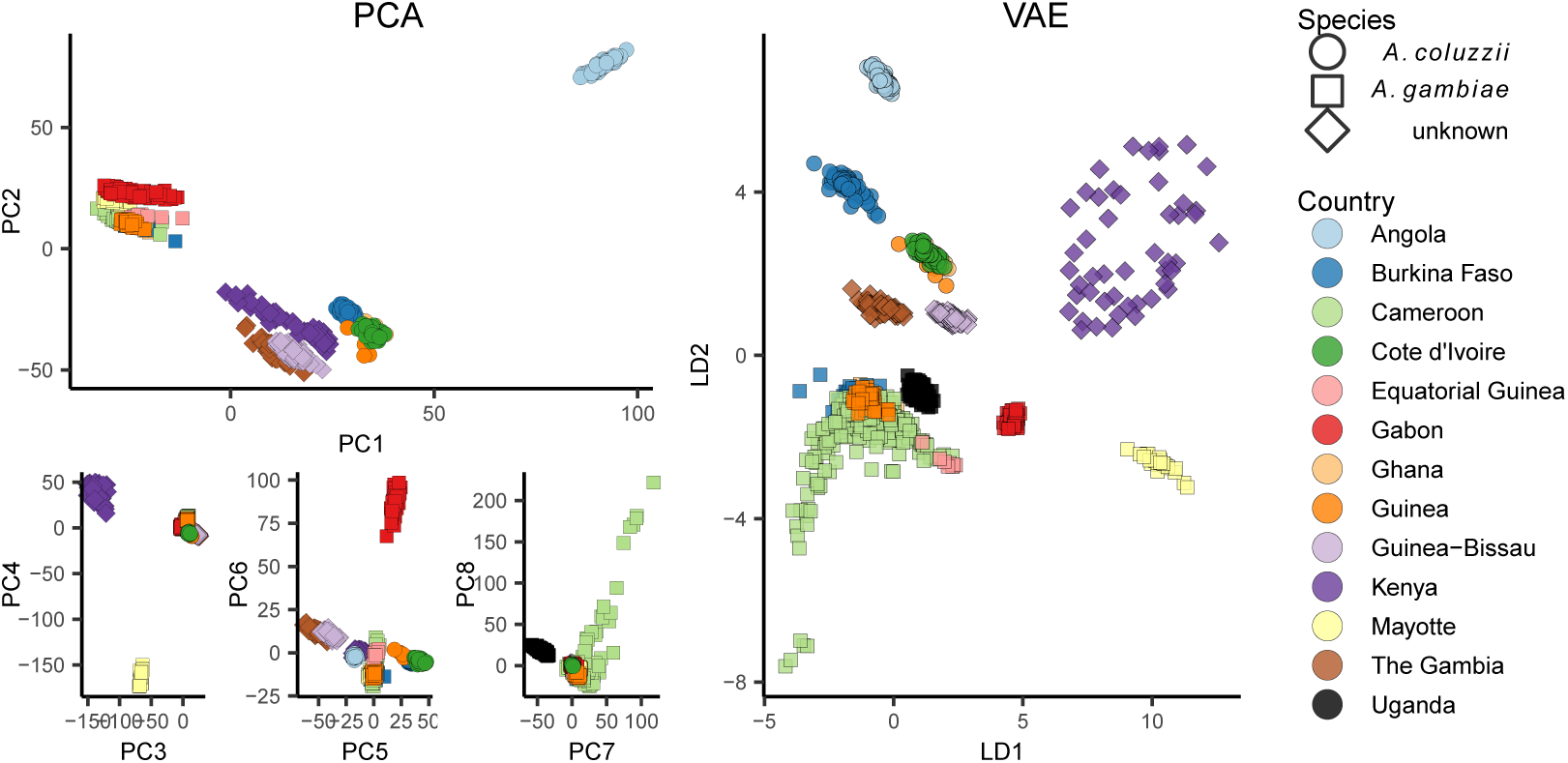
PCA (left) and VAE (right) run on 100,000 random SNPs from chromosome 3R of the AG1000G phase 2 data. Axes are flipped to approximate geography.

*A. gambiae / coluzzii* genomes are characterized by a series of well-studied inversions on chromosomes 2L and 2R (Coluzzi et al., 2002) which segregate within all populations and are associated with both malaria susceptibility and ecological niche variation (Riehle et al., 2017). The large 2La inversion contains at least one locus for insecticide resistance (*Rdl*), and has experienced multiple hard sweeps and introgression events in its recent history (Grau-Bové et al., 2020). Inversions have significant effects on local PCA (Li and Ralph, 2019) which often lead to samples clustering by inversion karyotype rather than geography on the first two PC axes (Ma and Amos, 2012).

To test how our VAE responds to inversions we fit models to SNPs extracted from 200,000 bp non-overlapping windows across the 2LA inversion in the AG1000G phase 2 data (Figure 6, Figure S11). We took an approach similar to Li and Ralph (2019) to summarize differences in latent spaces across windows while accounting for axis rotation and scaling. Latent dimensions were first scaled to 0 - 1 and the pairwise Euclidean distance matrix among individuals was calculated for each window to generate rotation- and scale-invariant representations of the latent space. We then calculated Euclidean distances among all pairs of per-window distance matrices, giving us a matrix representing relative differences in latent spaces across windows. Last, we used multi-dimensional scaling to compress this distance matrix to a single dimension, and plotted this value against genomic position across the 2La inversion region.

**Figure 6:**
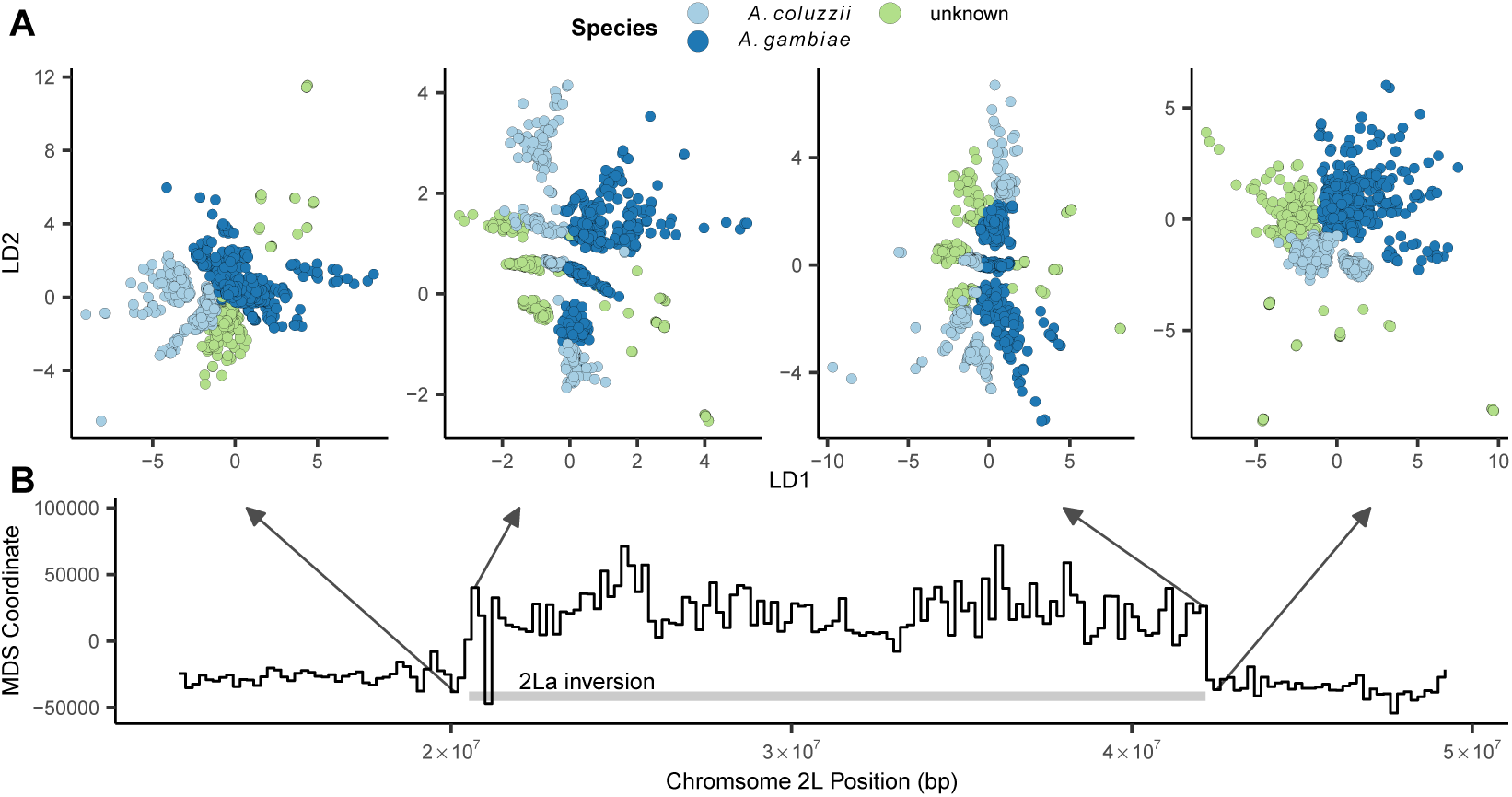
Latent spaces reflect inversion karyotypes at the 2La inversion in *A. gambiae / coluzzii*. A: VAE latent spaces for AG1000G phase 2 samples from windows near the 2La inversion breakpoints, colored by species. B: Multi-dimensional scaling values showing difference in the relative position of individuals in latent space across windows – high values reflect windows in which samples cluster by inversion karyotype, and low values by species.

This analysis found two clear classes of latent spaces inside and outside the inversion (Figure 6). Outside the inversion samples generally cluster by species and geography, while inside the inversion samples form three clusters corresponding to the homozygous and heterozygous inversion karyotypes, similar to results found with PCA (Grau-Bové et al., 2020; Riehle et al., 2017). Interestingly the VAE retains geographic and species clustering within inversion classes, but loads these aspects of structure on a different latent dimension than the karyotype clusters (e.g. LD1 reflects species clusters while LD2 reflects inversion karyotypes in the windows shown in Figure 6). Unlike PCA, latent dimensions from a VAE are not ranked by variance explained and nothing in the loss function incentivizes splitting particular aspects of variation onto separate axes, so we found this pattern of partitioning geographic and karyotypic signals somewhat surprising.

### Simulations and Sensitivity Tests

In general a method’s ability to detect population structure in a sample of genotypes scales with the degree of differentiation and the size of the genotype matrix. Patterson et al. (2006) found that there is a “phase change” phenomenon by which methods like PCA transition from showing no evidence of structure to strong evidence of structure when 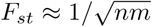, where *n* is the number of genotyped SNPs and *m* is the number of sampled individuals.

To compare the performance of PCA and VAE around this threshold we ran a series of two-population, isolation with migration model coalescent simulations in msprime (Kelleher et al., 2016) while varying the symmetric migration rate to produce an expected equilibrium *F*_*st*_ ranging from 0.0001 to 0.05. We sampled 50 diploid genomes from each population and downsampled the resulting genotype matrix to 10,000 SNPs. Given this sample size we expect the threshold for detecting structure to be approximately *F*_*st*_ = 0.001.

With tuned hyperparameters the VAE appeared slightly more sensitive to weak structure than the first two axes of a PCA (Figure 7). Both `popvae` and PCA reflect some population structure at *F*_*st*_ >= 0.005 (though this is clearer in the VAE) but none at *F*_*st*_ <= 0.001, consistent with Patterson et al. (2006)’s “phase change” suggestion. However the VAE’s performance was highly sensitive to hyperparameter tuning on this dataset. At default settings `popvae` latent spaces reflect no clear structure until *F*_*st*_ = 0.05 (Figure S12,Figure S13). In particular we found that increasing the ‘`patience`’ parameter to 500 was necessary for even marginal performance in this case, and running a grid search across network sizes was needed to match PCA’s sensitivity to weak structure.

**Figure 7:**
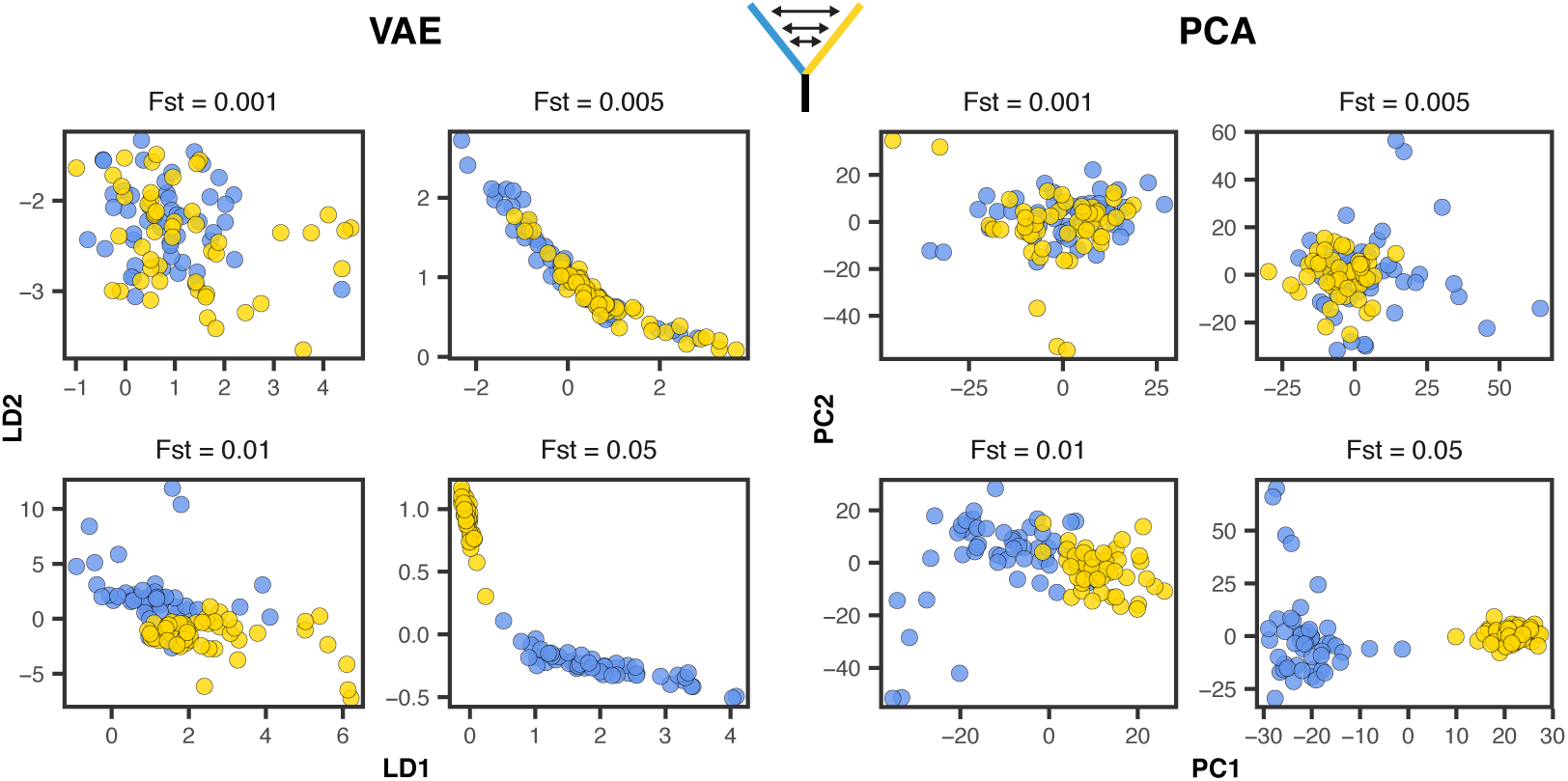
VAE latent spaces and PCA run on two-population coalescent simulations with *F*_*st*_ varying from 0.0001 − 0.05. Points are colored by population. `popvae` was run with tuned hyperparameters and patience set to 500. See Figure S12 for (much worse) performance with default settings.

### Comparison with UMAP and t-SNE

In addition to PCA we also compared the VAE’s latent spaces to t-SNE (Maaten and Hinton, 2008) and UMAP (Diaz-Papkovich et al., 2019) (Figure S14, Figure S15), both of which have been used recently for population genetic visualization. We first ran both methods on the top 15 PC axes (following Diaz-Papkovich et al. (2019)) with default settings on the human and *Anopheles* datasets and used the R packages ‘umap’ (Konopka, 2019) and ‘tsne’ (Donaldson, 2016) as our reference implementations.

For HGDP data both UMAP and t-SNE produce latent spaces that roughly correspond to continental regions (Figure S14). Running both methods at default settings, UMAP’s latent space was much more tightly clustered – for example grouping all samples from Africa into a single small region. Similar patterns were seen in the AG1000G data (Figure S15) – both t-SNE and UMAP produce latent spaces that strongly cluster sample localities and species. However, global geometry appeared to be poorly preserved in t-SNE and UMAP latent spaces. That is, though clusters in latent space correspond to sampling localities, distances among clusters do not appear to meaningfully reflect geography or genetic differentiation.

To compare how well different methods reflect geography we compared pairwise distances among individuals in latent and geographic space for Eurasian human samples (HGDP regions Europe, Central/South Asia, the Middle East, and East Asia). Geographic distances were great-circle distance calculated on a WGS84 ellipse with the R package ‘sp’ (Pebesma et al., 2012). Distances were scaled to 0-1 for this analysis, and we calculated the coefficient of determination (*R*_2_) across geographic and latent-space distance for each method as a metric. VAE latent space distances have the strongest correlation with geographic distance (Figure 8; *R*_2_ = 0.659), followed by PCA (*R*_2_ = 0.561), UMAP (*R*_2_ = 0.529), and t-SNE (*R*_2_ = 0.342).

**Figure 8:**
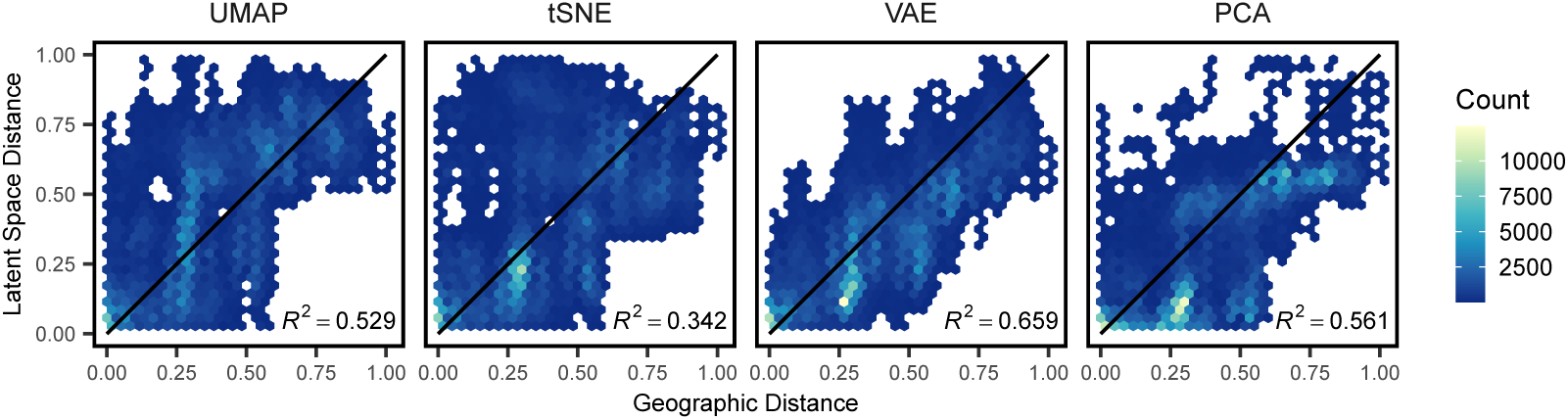
Comparing pairwise distances in geographic and latent space for Eurasian human genotypes across four dimensionality reduction methods run at default settings. All distances are scaled to 0-1. Black lines show a 1:1 relationship.

Finally to test how parameter tuning of tSNE and UMAP impacts our results, we reproduced our analysis of HGDP data using double and triple the default values for `n neighbors` (UMAP) and `perplexity` (tSNE). Though scatter plots are visually similar at these settings (Figure S16) the correlation between latent-space and geographic distances of Eurasian samples is improved in both methods at double default settings (t-SNE: *R*_2_ = 0.631, UMAP: *R*_2_ = 0.611; Figure S17). At triple default settings we observed slightly better performance for tSNE and slightly worse for UMAP (Figure S18, Figure S19).

### Run Times and Computational Resources

We compared `popvae`’s run times to PCA, UMAP, and t-SNE using sets of 100,000 and 10,000 SNPs from the HGDP as described above. `popvae` was run using default settings (i.e. fitting a single network rather than running a grid search over network sizes) using a consumer GPU (Nvidia GeForce RTX 2070). PCAs were run in the python package scikit-allel (Miles and Harding, 2017), which in turn relies on singular-value decomposition functions from the numpy library (Oliphant, 2006–).

`popvae` was much slower than PCA or UMAP, but comparable to running t-SNE on PC coordinates. However for datasets of the size we tested here none of these run times present significant challenges – all methods return sample latent coordinates in less than five minutes. We have not conducted exhaustive tests on CPU training times for `popvae`, but in general find these to require at least twice as much time as GPU runs.

However for larger datasets we expect `popvae`’s run time performance would suffer further in comparison to PCA and UMAP. The major computational bottleneck is loading tensors holding weights for the input and output layers of the encoder and decoder networks into GPU memory. These tensors have dimensions `n snps x network width` so they become extremely large when running on large genotype matrices. Our development machine has 8GB GPU RAM and can process up to roughly 700,000 SNPs in a single analysis using a 128-unit-wide network. Throughout this study we have limited our analysis to relatively small subsets of genome-wide SNPs to allow us to explore a range of network sizes in reasonable time. Scaling up to a single model fit to all genome-wide SNPs – on the order of 107 for datasets like the HGDP – would require access to specialized hardware with very large GPU memory pools.

**Table 1:** Run times for VAE, PCA, UMAP, and t-SNE HGDP data. UMAP and t-SNE were run on the top 20 PC axes (run times thus include running the PCA).

| run time (s) | SNPs | method |
| --- | --- | --- |
| 204.4 | 100,000 | VAE |
| 3.6 |  | PCA |
| 6 |  | UMAP |
| 124.8 |  | t-SNE |
| 78.8 | 10,000 | VAE |
| 0.5 |  | PCA |
| 2.7 |  | UMAP |
| 119.5 |  | t-SNE |

### Generating Genotypes

The VAE framework also allows us to generate genotypes characteristic of a given population by sampling from the latent space of a trained model. Simulated genotypes generated by process-based models like the coalescent are a key tool in population genetics, because they allow us to explore the impact of various generative processes – demography, selection, etc – on observed genetic variation (Adrion et al., 2020a). In contrast `popvae`’s generative model provides essentially no mechanistic insight beyond the strong observed correlation of latent and geographic spaces. However, if the VAE accurately reproduces characteristics of real genotypes it could be a fast alternative to simulation that does not require parameterizing a custom demographic model.

We compared these approaches by analyzing empirical data from European (CEU), Han (CHB), and Yaruban (YRI) human genotypes in the 1000 Genomes Project data (Consortium et al., 2015). We first subset 50 samples from each population and then fit a 2-dimensional `popvae` model to all SNPs from chromosome 22. To generate genotypes we drew a sample from the latent distribution of each individual and passed these coordinates to the trained decoder network. We interpret the sigmoid activation output of our decoder as the probability of observing a derived allele at each site, and generate derived allele counts by taking two draws from a binomial distribution with *p* = *g*_*i,j*_ where *g*_*i,j*_ is the decoder output for individual *i* at site *j*.

As a baseline comparison we used coalescent simulations from the `standardpopsim` library (Adrion et al., 2020a) of the 3-population out-of-Africa model (`OutOfAfrica 3G09`) – a rigorously tested implementation of the demographic model fit to the joint site frequency spectrum in Gutenkunst et al. (2009) using the `msprime` coalescent simulator (Kelleher et al., 2016). For this comparison we changed `standardpopsim`’s default human mutation rate of 1.29 × 10^−8^ to 2.35 × 10^−8^ to match the rate used in Gutenkunst et al. (2009), used the `HapMapII GRCh37` recombination map for chromosome 22, and sampled 100 haploid chromosomes from each population.

Last, we examined three facets of population genetic variation on real, VAE-generated, and simulated genotype matrices: the site frequency spectrum, the decay of linkage disequilibrium with distance along the chromosome, and embeddings from a PCA. These analyses were conducted in scikit-allel (Miles and Harding, 2017) after masking genotypes to retain only sites with the most stringent site accessibility filter (“P”) in the 1000 genome project’s phase 3 site accessibility masks. LD statistics were calculated only for YRI samples using SNPs between positions 2.5 × 10^7^ and 2.6 10^7^ in the hg18 reference genome and summarized by calculating the mean LD for all pairs of alleles in 25 distance bins (similar results in three different genomic windows are shown in figure S20). Results are plotted in figure 9. In general we found all methods produce similar results in a plot of the first two PC axes, suggesting they capture broad patterns of allele frequency variation created by population structure. The site frequency spectrum is also very similar for the VAE and real data, while the simulated genotypes suffer from a scaling issue. This could reflect differences in the input data – Gutenkunst et al. (2009) fit models to an SFS calculated from a set of sanger-sequenced loci in 1000 genomes samples, rather than the short-read resequenced SNPs from the 1000 Genomes project we use – or an inaccuracy in one of the constants used to convert scaled demographic model parameters to real values (accessible genome size, generation time, or mutation rate). LD decay shows the largest difference among methods. Simulation and real data both reflect higher LD among nearby SNPs which decays with distance, while the VAE genotypes produced no correlation between distance along a chromosome and pairwise LD.

**Figure 9:**
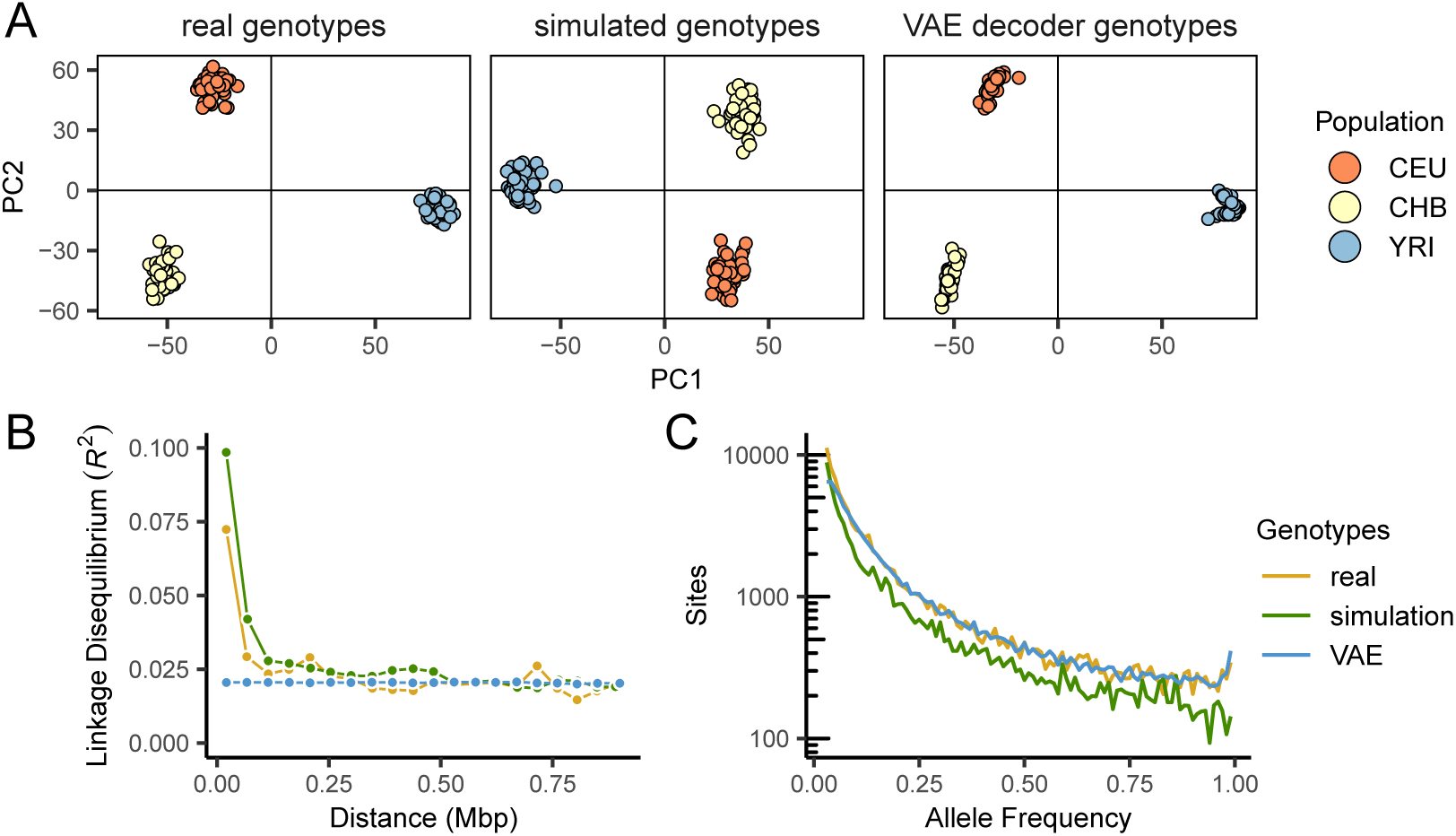
Comparing real, VAE-generated, and simulated genotype matrices for three populations from the 1000 genomes project. The VAE decoder and coalescent simulation produce similar results in genotype PCA (A), but the VAE fails to reproduce the decay of LD with distance along the chromosome seen in real data (B). The site frequency spectrum is very similar for real and VAE-generated genotypes, but suffers from scaling issues in the coalescent simulation (C).

These differences reflect the strengths and weaknesses of each method. The VAE decoder doesn’t require a pre-defined demographic model and by design exactly fits the matrix size of input empirical data, so it should not suffer from the scaling issues that frequently impact population genetic models. But the lack of mechanistic biological knowledge in its design means it misses obvious and important features of real sequence data like the decline of LD with distance. In this case the lack of LD decay in VAE decoder sequences means this implementation should not be used for testing properties of analyses like GWAS, in which LD among a subset of sequenced loci and an unknown number of truly causal loci is a crucial parameter. Though other network designs (e.g. a convolutional neural network Flagel et al. (2019) or a recurrent neural network Adrion et al. (2020b)) could potentially address the specific shortcoming of LD decay, the general problem of a non-mechanistic generator failing to mimic features of the data produced by well-understood processes seems intrinsic to the machine learning approach.

## Discussion

Dimensionality reduction of genotypic variation is a key analytic tool in modern genomics and their visualizations are often the central figure of a genetic study. For example, Antonio et al. (2019) studied a 10,000-year transect of genotypes from Rome and extensively used PCA to visualize changes in ancestry in the city over time. In cases like this producing informative plots of population structure is a requisite step for the analysis and can shape the way data is interpreted both by authors and readers.

In this study we demonstrate how variational autoencoders can be used for visualization and low dimensional summaries of genotype data. Variational autoencoders have at least two attractive properties for genetic data: they allow users to define the output dimensionality, and they preserve global geometry (i.e., relative positions in latent space) better than competing methods. As we have shown in humans and mosquitoes, this allows users to generate visualizations that summarize relationships among samples without either comparing across several panels (as with PCA) or attempting to ignore possibly spurious patterns of global structure (as with t-SNE and UMAP).

An additional attractive property of VAEs is that they are generative models. That is to say that VAEs allow us to create genotypes that capture aspects of population genetic variation characteristic of the training set. This is done by taking samples from the estimated latent space and passing forward into data space. Though in theory this could be used as an alternative to simulation, our implementation fails to replicate at least one important aspect of real genomes – the decay of linkage disequilibrium with distance along a chromosome – and thus offers limited utility for tasks such as boosting GWAS sample sizes or as a substitute for simulation. We point researchers interested in generating genotypes via deep learning approaches to recent work by Yelmen et al. (2019) and Montserrat et al. (2019), which describe similar, deep learning based methods more tightly aimed at generating realistic genotypes.

The are also several significant limitations of our method as a visualization tool. One issue is that we lack a principled understanding of how the VAE output maps to parameters of idealized population models like the coalescent (Kingman, 1982). This is in contrast to PCA, which was first applied to genetic data with little theoretical background (Menozzi et al., 1978) but is now fairly well characterized in reference to population genetic models (McVean, 2009; Novembre and Stephens, 2008).

Hyperparameter tuning is another challenge. As we showed, `popvae` has many hyperparameters that significantly affect the output latent space and no principled way to set them *a priori*. Though we include a grid-search function for network sizes, this is slow and is still dependent on other hyperparameters – like the patience used for early stopping, or the learning rate of the optimizer – which we have set to defaults that may not be optimal for all datasets. This is not a unique issue to VAEs; opaque hyperperameters of methods like t-SNE and UMAP can significantly affect embeddings (Kobak and Linderman, 2019), and preprocessing choices such as how to scale allele counts prior to PCA dramatically vary the appearance of final plots (Patterson et al., 2006). However it does require extra work on the part of users interested in exploring the full parameter space.

A parallel issue is stochasticity in the output. Stochasticity is introduced by the random test/train split, parameter initialization states, and even the execution order of operations run in parallel on GPU during model training. Though all but the last of these can be fixed by setting a random seed, which itself could be (unfortunately) seen as a hyperparameter, there is no obvious way to compare models fit to different validation sets in a world of limited training examples. This introduces noise which could potentially allow users to cherry-pick a preferred latent space.

For example, one run of our best-performing network architecture on the HGDP data produced a latent space with in which samples Papua New Guinea and Bougainville are separated by roughly the same distance as samples from north Africa and East Asia. In contrast all other fits of the same network architecture cluster these samples (Figure S2, see the top middle panel). We chose a latent space for the main text that lacked this feature because it occurred in only one training run, but acknowledge this procedure is sub-optimal. Developing a method to summarize across multiple latent spaces, perhaps via ensemble learning approaches, would be useful for postprocessing VAE output when latent spaces vary.

The last major shortcoming is computational effort. `popvae` is much slower and more computationally intensive than PCA, and requires specialized and expensive GPU or TPU hardware to run on large sets of SNPs. Future developments in both hardware and software will likely alleviate this issue somewhat, but at present it may make the method difficult to apply to the increasingly common whole genome resequencing data now being generated for many species.

One important question we did not explore in this study is whether VAE latent space co-ordinates offer any improvement over PCA when used as covariates to correct for population structure in GWAS (Price et al., 2006). UMAP and t-SNE are generally thought to be inappropriate for this use because of their failure to preserve global geometry (Diaz-Papkovich et al., 2019), but because the VAE appears to strongly reflect geography in humans it may be useful for this task. Testing this aspect of the VAE could be done in simulation but would benefit from empirical investigations in large human datasets – a task which is beyond the scope of the present study, but perhaps fruitful for further investigation.

Here we have shown that our implementation of a VAE, `popvae`, can produce informative visualizations of population genetic variation and offers some benefits relative to competing methods. However our approach is just one implementation of a huge class of potential models falling under the VAE umbrella. Altering the prior on the latent space (Davidson et al., 2018), the weighting of the loss function (Higgins et al., 2017), or the type of neural network used in either the encoder or decoder all offer avenues for further research and potential improvement (see also Appendix 1, where we briefly describe alternate approaches we experimented with). Entirely different methods of visualizing population structure which focus on genetic variants rather than individuals, like that proposed in Biddanda et al. (2020), also offer a complementary perspective on the nature of genetic differentiation. As population genetic data becomes increasingly common across evolutionary biology we anticipate visualization techniques will receive increased attention from researchers in many areas, and believe VAEs offer a promising avenue for research.

## Acknowledgements

We thank Peter Ralph for the suggestion to use binomial sampling to bin the decoder output, and other members of the Kern-Ralph co-lab for comments on the software and manuscript. Thanks to the reviewers and editors for your comments. Audry Gill developed a successful pilot of this project early after its inception. CJB was supported by NIH awards R01GM117241 and F32GM136123. ADK was supported under NIH awards R01GM117241 and R01HG010774.

## Appendix 1

### Other Things We Tried That Didn’t Work

We tried a bunch of things while developing `popvae`. Here we document some of our dead-ends in the hope they may be useful to others developing similar methods.

#### 0.0.1 A Convolutional Neural Network

We first developed `popvae` using convolutional neural networks (CNNs) for both the encoder and decoder. The feed-forward network we use here was originally intended as a naive baseline for comparing our CNN performance, but it turned out to be faster and more accurate (that is, lower validation loss), and had much lower memory requirements than any CNN we tried. These included 2D CNNs run on phased haplotypes, 1D CNNs run on unphased genotype counts, hybrid CNN+feed-forward networks stacking convolutional and dense layers in succession, and restricting the CNN to either the encoder or the decoder.

#### 0.0.2 A Recurrent Neural Network

We also tested recurrent neural networks (using the `cudnnGRU()` layer in keras) as one or both of the encoder/decoder pair. Due to memory limitations we were only able to test relatively small, shallow networks with this approach (width 32, depth up to 3). Like the CNNs these were slower, less accurate, and more resource-intensive than the dense network we describe in the main text.

#### 0.0.3 Skipping the Validation Set

It would be nice to not need a validation set. The train/test split introduces extra stochasticity and you have to ignore some hard-earned data in training.

Unfortunately we couldn’t find a good way of setting the learning rate scheduler or establishing a good stopping time for model training without a validation set. Training on all samples leads to constantly decreasing loss so all training runs go to the maximum number of epochs. Examining the progress of latent spaces through model training for these runs, the encoder seems to quickly identify and then refine structure in the input samples, but eventually samples begin to cluster in a ring around the origin at 0,0. This appears to reflect the Gaussian prior on the latent space dominating the loss function as the reconstruction error approaches some lower bound. In runs with validation sets we observed that validation loss typically increases once points begin circling the origin (Figure S1), suggesting it reflects a typical overfitting behavior. Unfortunately the number of epochs needed for this to occur is different for every dataset and we found no general solution for estimating its location other than a validation set.

So we recommend using a validation set of at least 10%, and comparing latent spaces from runs with multiple starting seeds (and so different train/validation splits). In a pinch, users can set `--train prop 1` to train on all samples and heuristically examine latent spaces output during training to figure out a good stopping point.

#### 0.0.4 Batch Normalization

Putting a batch normalization layer anywhere in either the encoder or decoder made validation loss worse in all of our tests.

#### 0.0.5 Dropout

As above, dropout layers either made no difference or yielded slightly higher validation losses no matter where we put it.

#### 0.0.6 Reweighting the Loss Function

Higgins et al. (2017) proposed a modification of the standard VAE loss function which amounts to multiplying the KL divergence by a factor *β*. This puts extra weight on the normal prior of the latent space and on the MNIST dataset delivered more clustered and interpretable latent spaces. Unfortunately the only suggested method for estimating *β* in a truly unsupervised setting like ours is heuristic examination of model output. We experimented with several values and found no consistent benefits either in latent space or validation loss relative to our baseline approach. However this seems like a productive area for further investigation and we plan to continue experimenting along these lines (and encourage others to do so as well).

## Supplementary Figures and Tables

**Figure S1:**
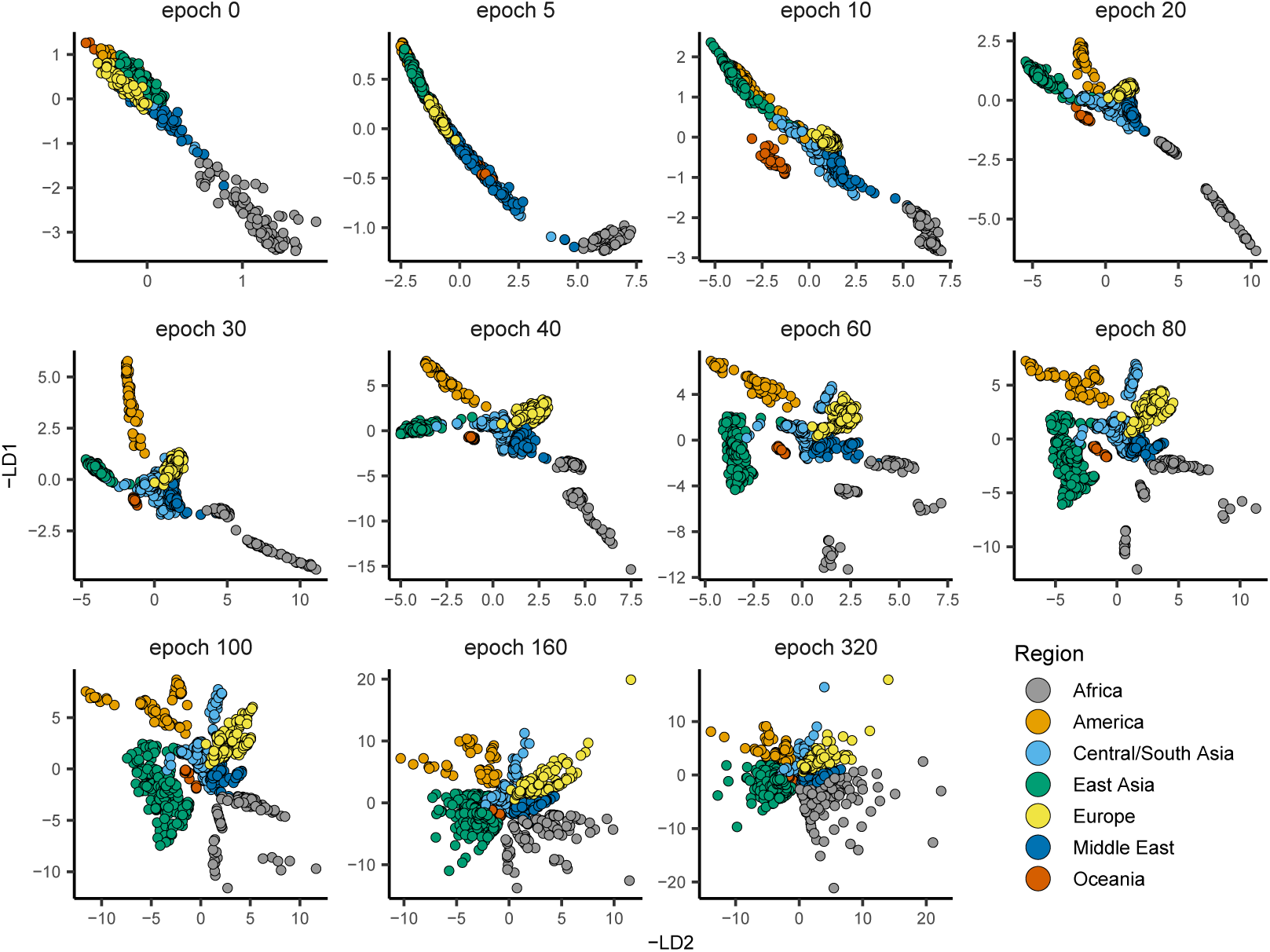
Latent spaces output during model training for HGDP data. Here `patience` was increased to 500 to show the overfitting behavior of `popvae`’s latent space. In this run validation loss was minimized at epoch 59.

**Figure S2:**
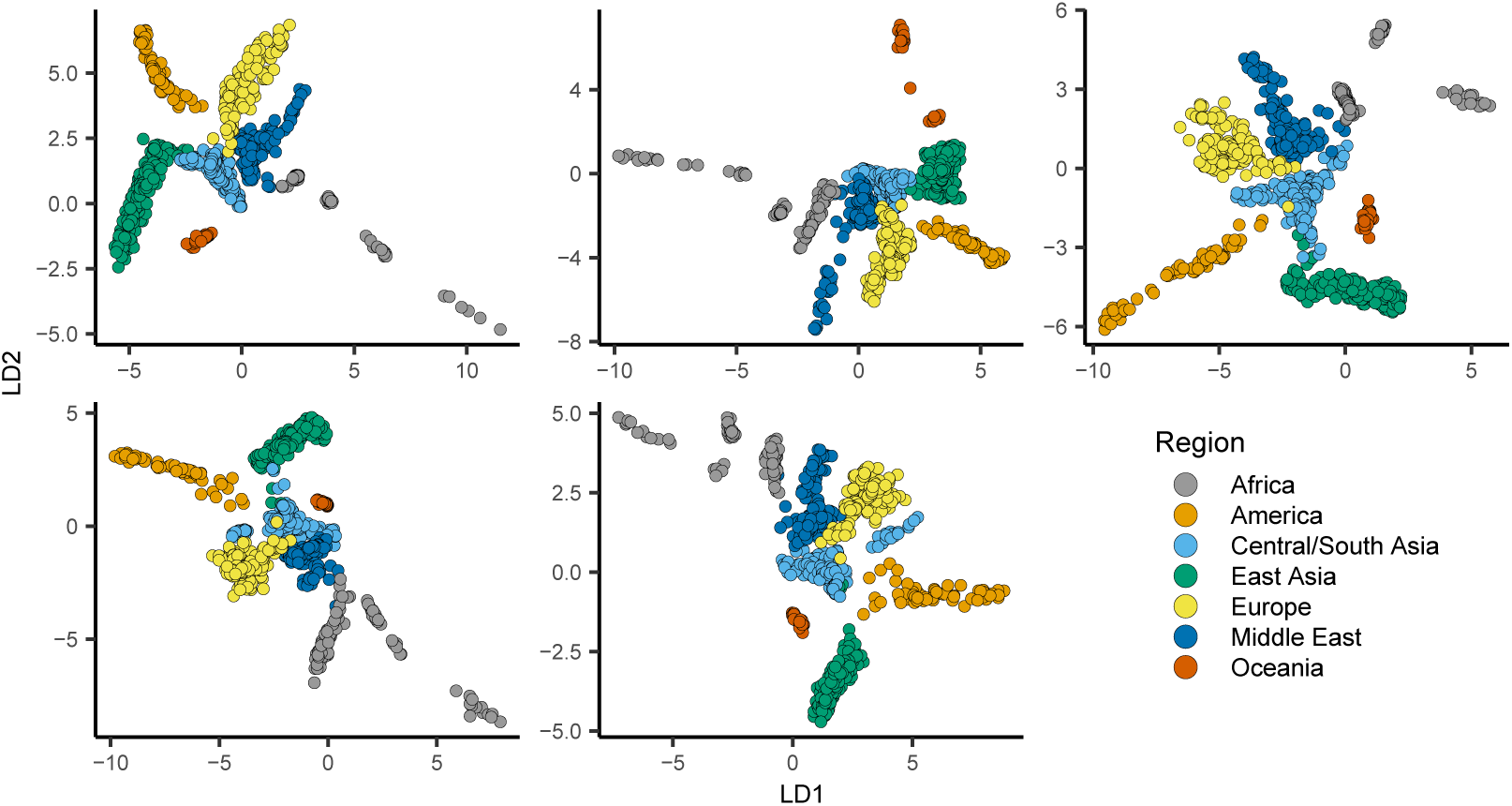
`popvae` latent spaces from runs with default hyperparameters and different random seeds.

**Figure S3:**
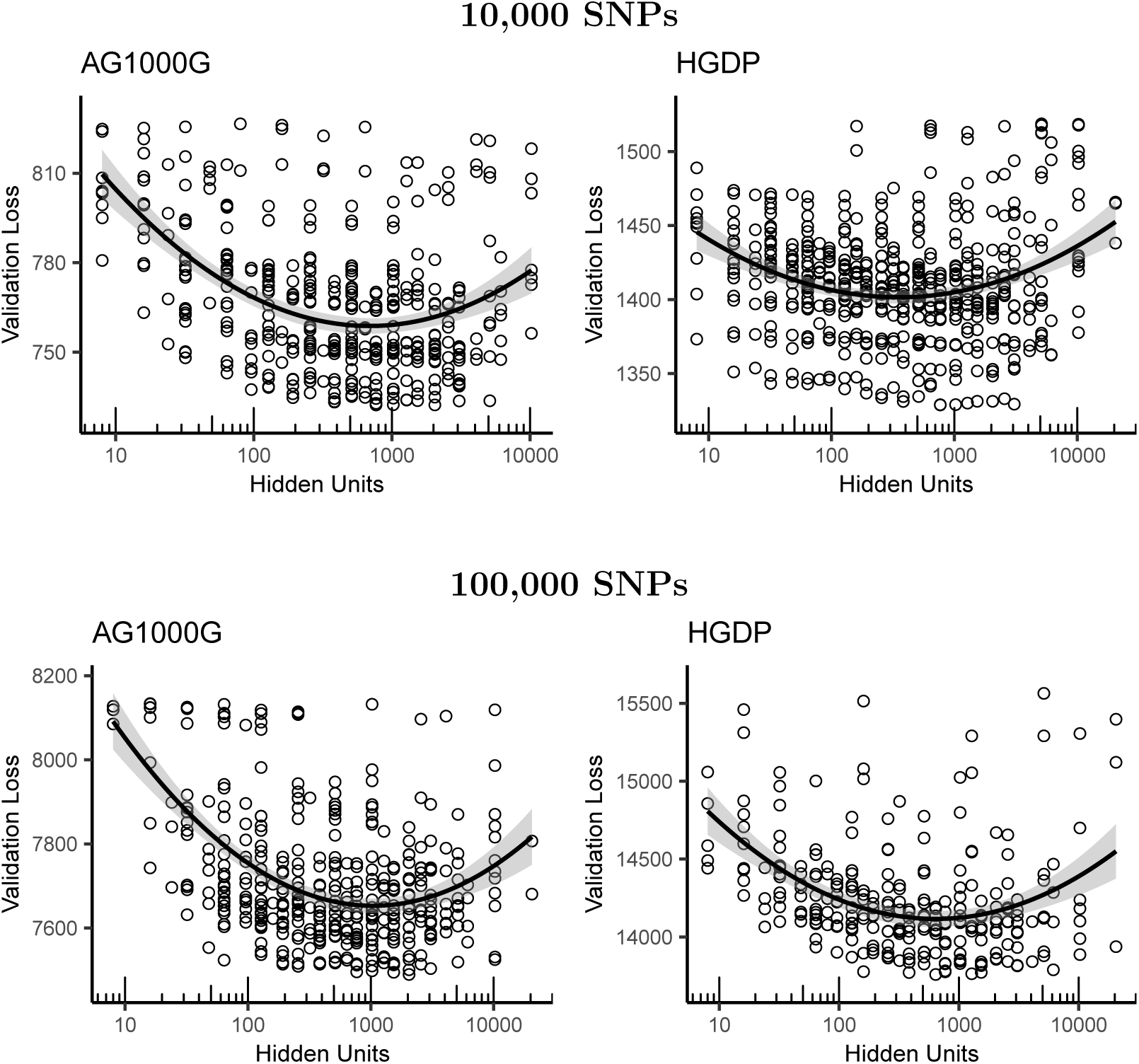
Validation loss as a function of the number of hidden units in a network for models fit to 100,000 SNPs selected randomly from *Anopheles* chromosome 3R and human chromosome 1. See table 1 for model rankings. Curves are quadratic least-squares model fits.

**Figure S4:**
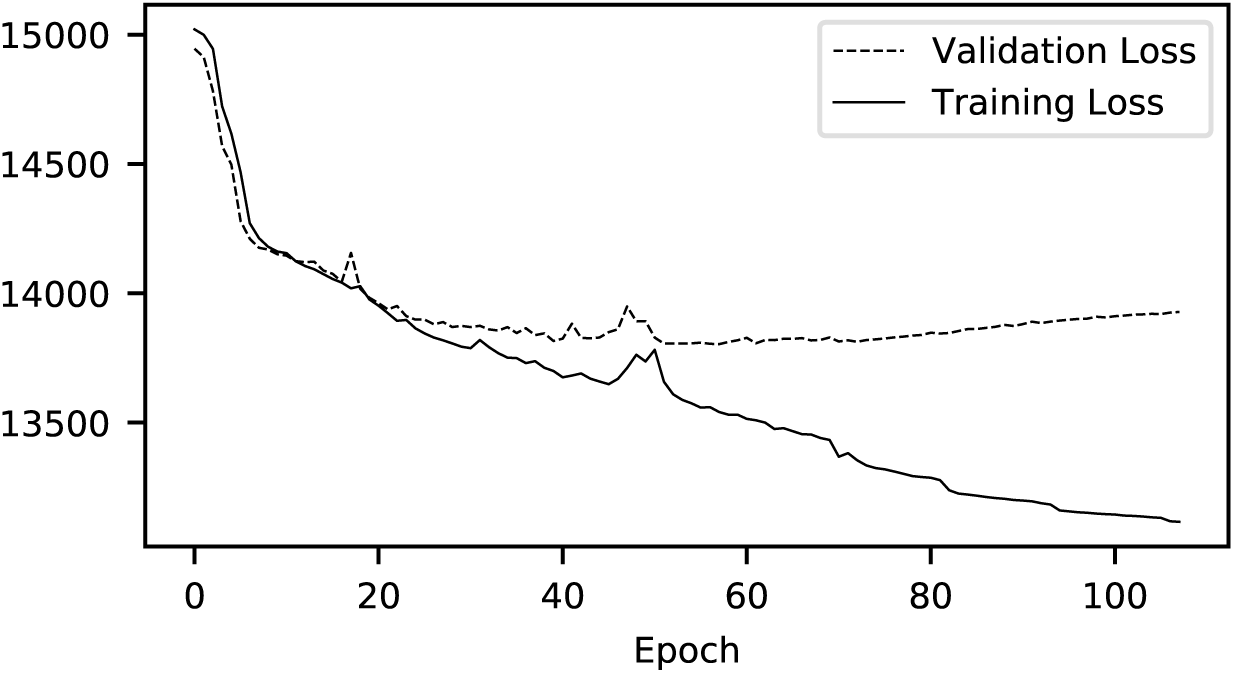
Example training history plot of showing training and validation loss by epoch during model training.

**Figure S5:**
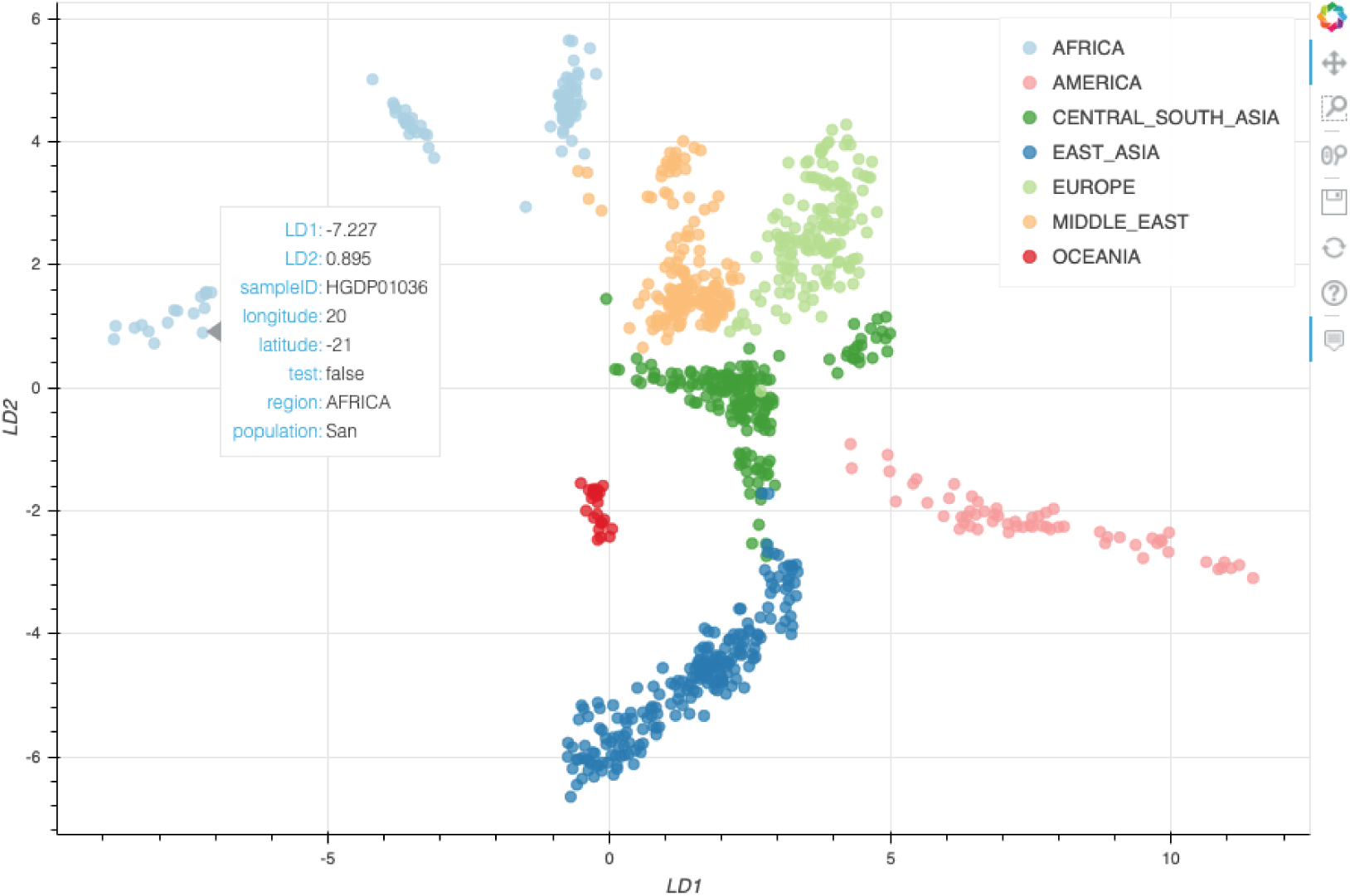
Example interactive plotting with scroll-over metadata.

**Figure S6:**
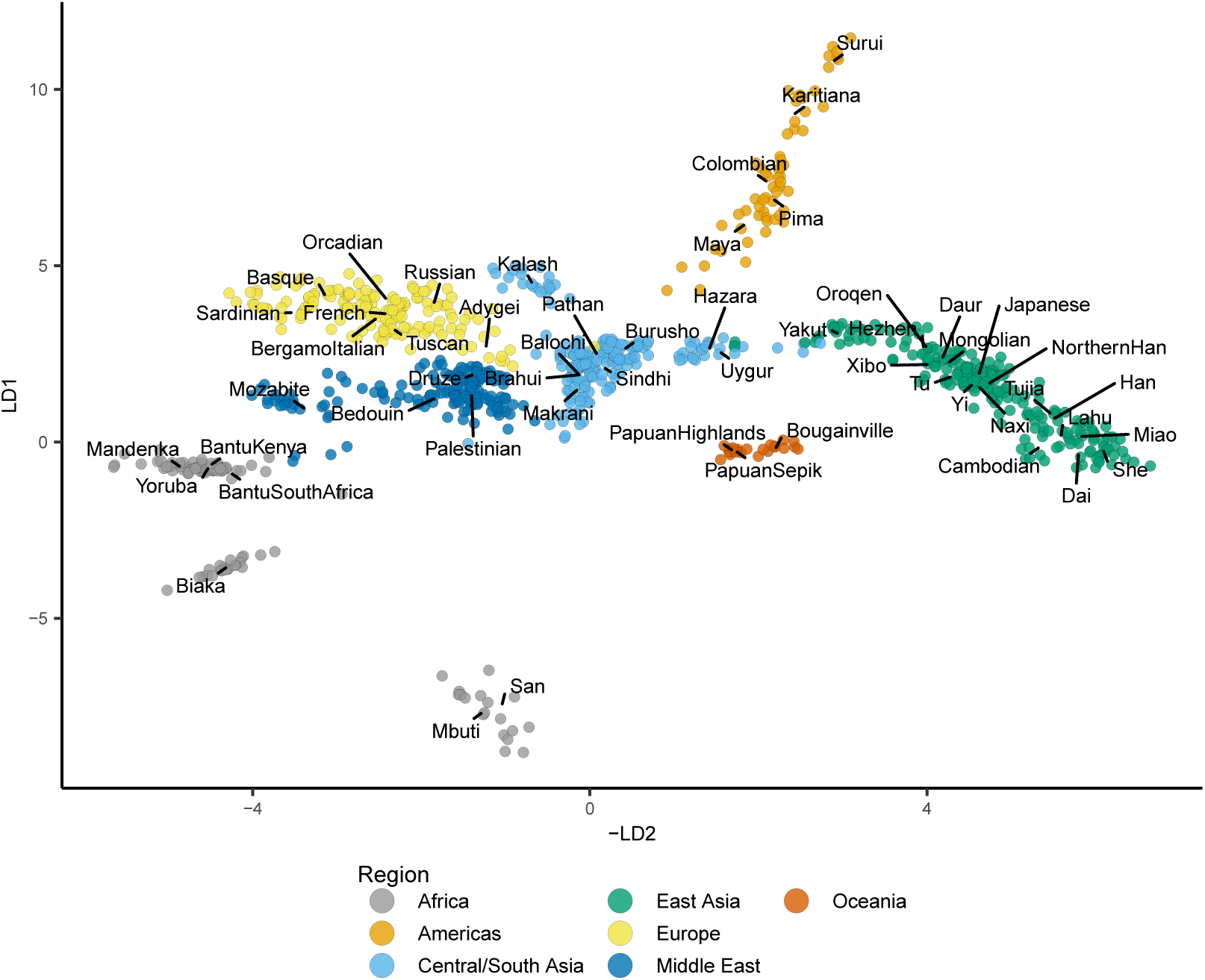
Latent space for 100,000 SNPs from chromosome 1 of the HGDP cohort (see Figure 2), with population centroids labeled.

**Figure S7:**
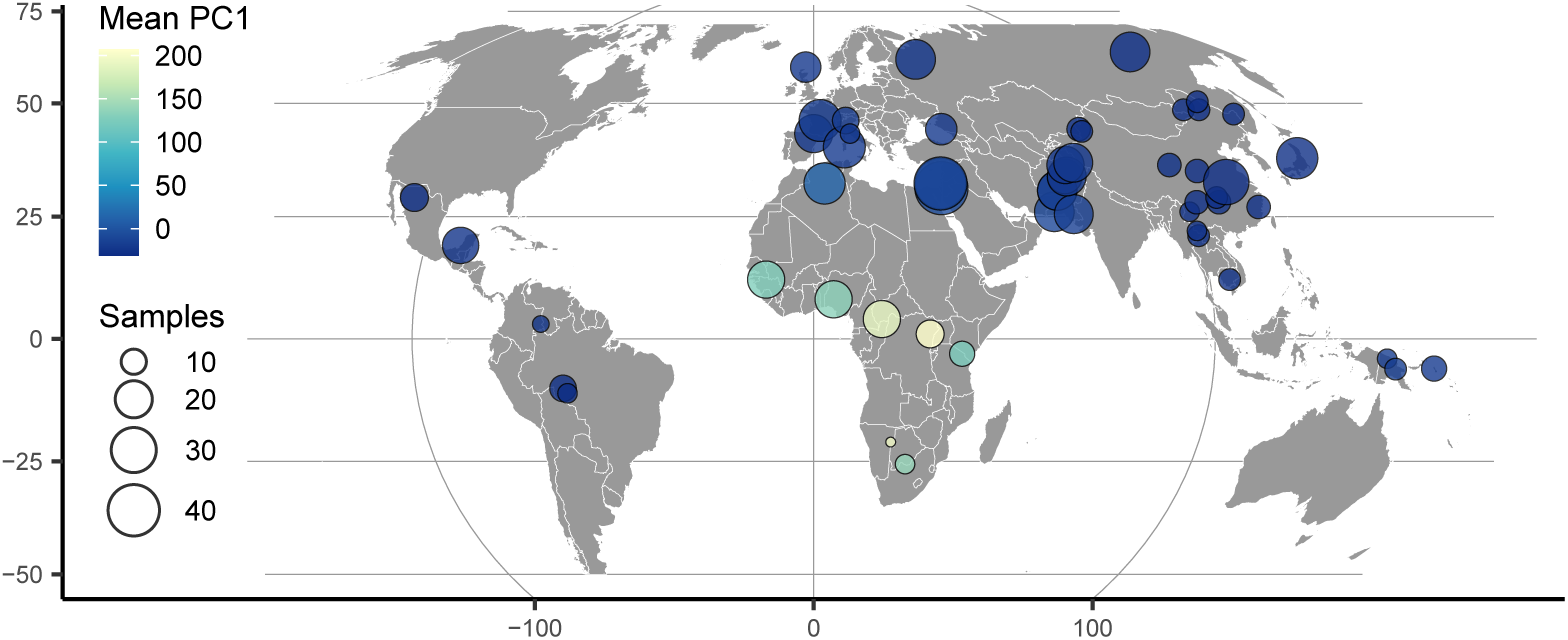
The first PC axis for HGDP SNPs summarized on a map as in Figure 3. Points show approximate population locations and are colored by the mean PC1 coordinate for each HGDP population. Densities show the distribution of PC1 scores for each HGDP region.

**Figure S8:**
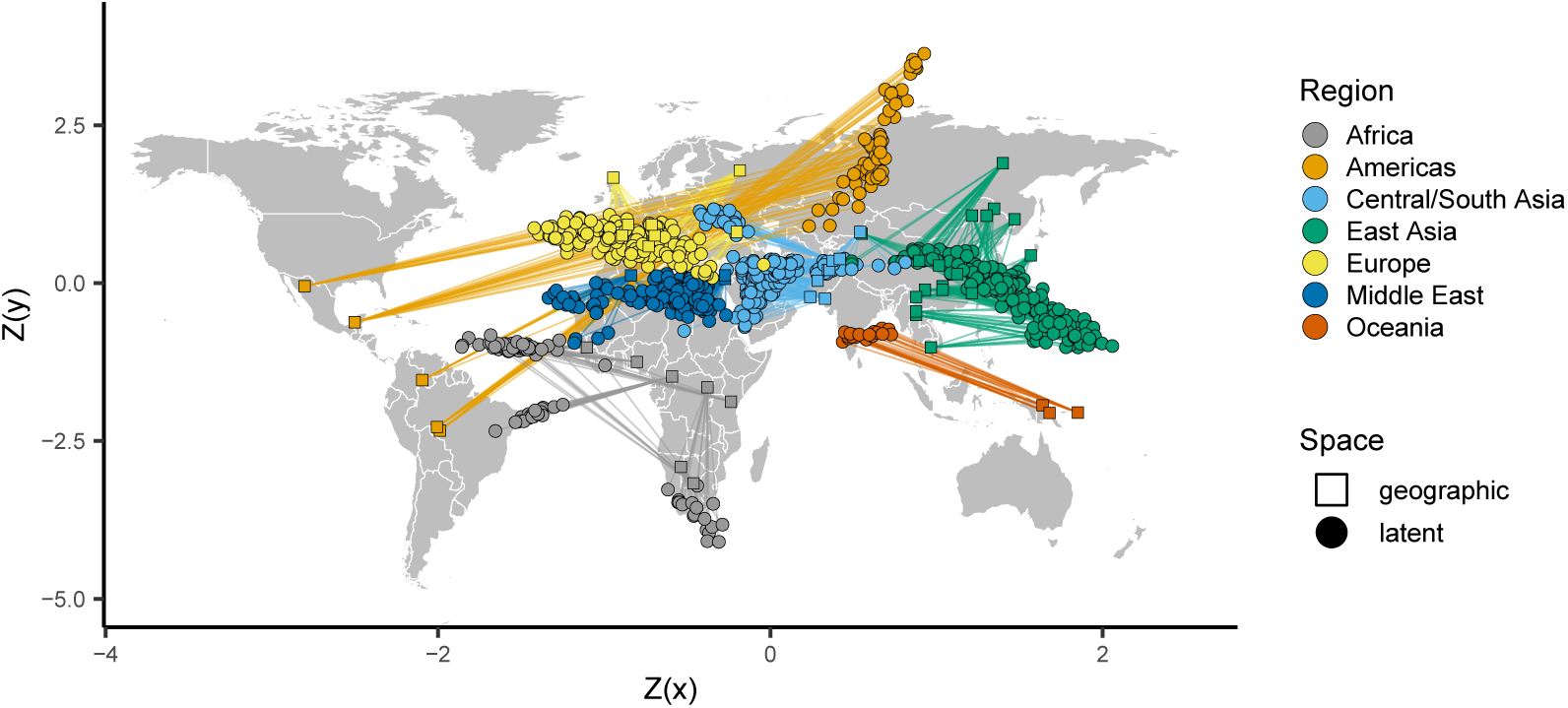
Comparing the VAE latent space with the geography of sampling localities HGDP samples. Circles show z-normalized sample locations in latent space and squares show the corresponding location in geographic space.

**Figure S9:**
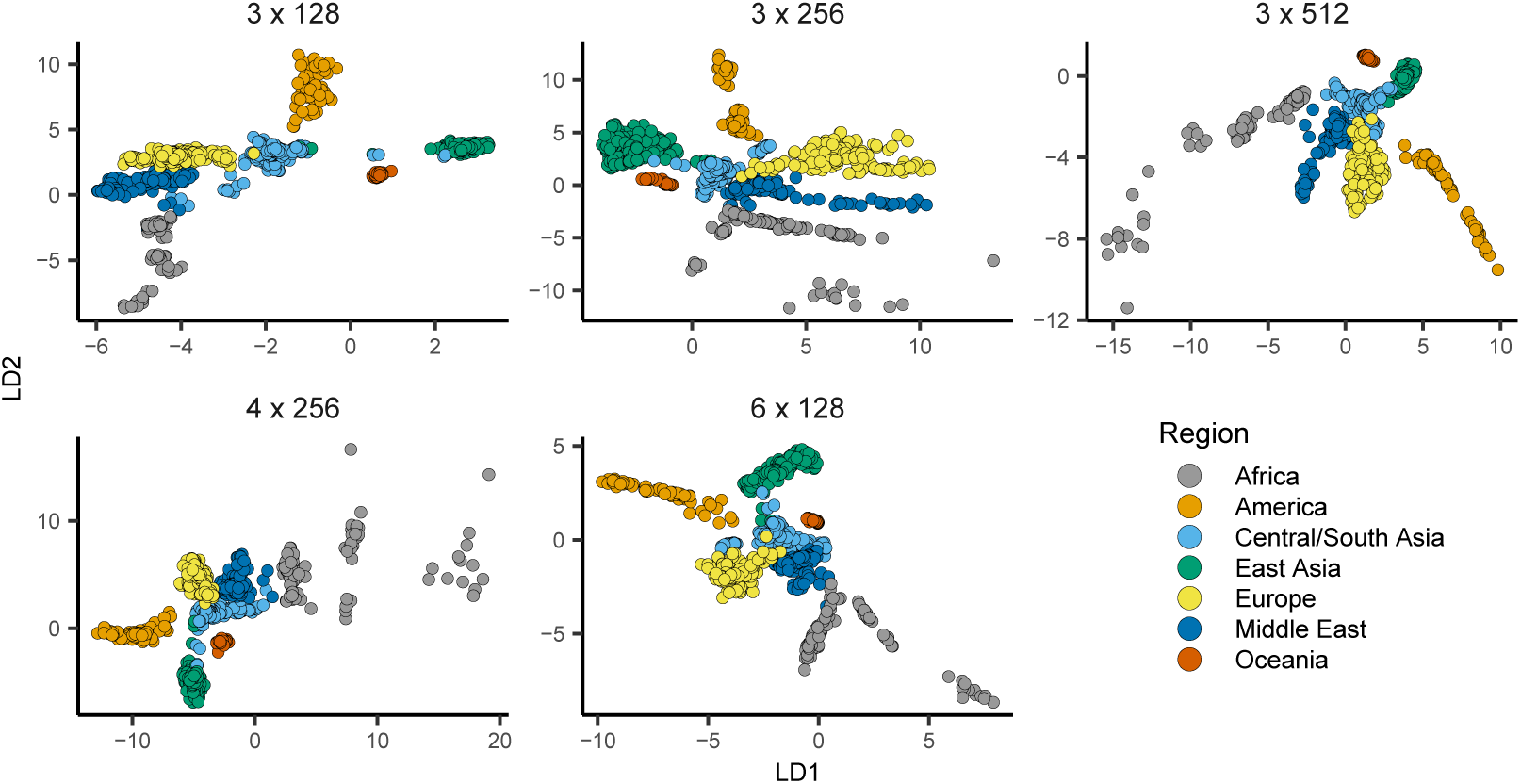
`popvae` latent spaces from runs with the same random seed and the top five network sizes by validation loss. Network sizes are listed as ‘`depth` x `width`’.

**Figure S10:**
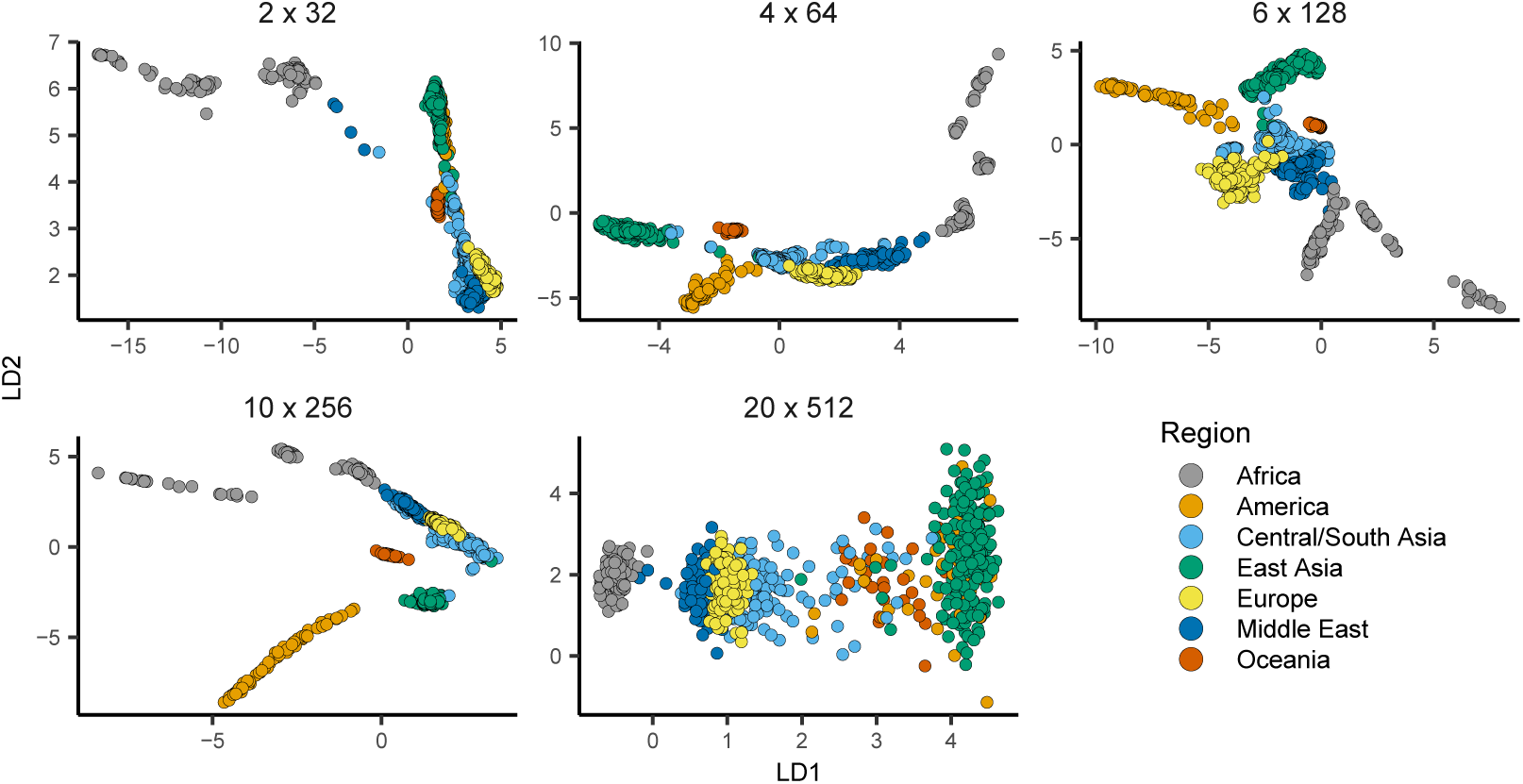
`popvae` latent spaces from models across the range of sizes tested. Network sizes are listed as ‘`depth` x `width`’.

**Figure S11:**
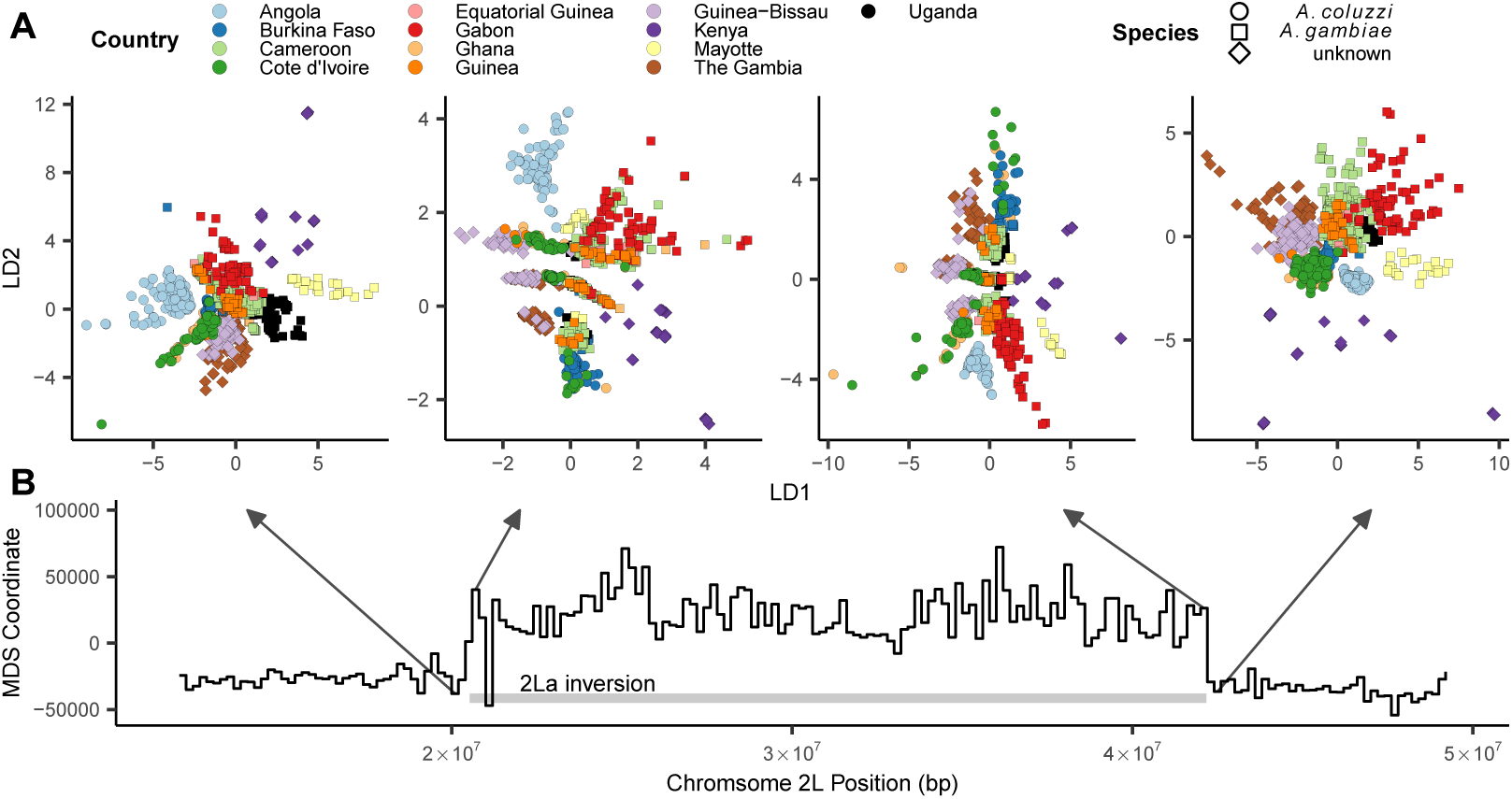
Latent spaces reflect inversion karyotypes at the 2La inversion in *A. gambiae / coluzzii*. A: VAE latent spaces for AG1000G phase 2 samples from windows near the 2La inversion breakpoints, with shapes indicating species and colors the country of origin. “Unknown” species localities include populations from Kenya, Guinea-Bissau, and the Gambia, for which diagnostic PCR markers are inconsistent or fail to amplify. B: Multi-dimensional scaling values showing difference in the relative position of individuals in latent space across windows – high values reflect windows in which samples cluster by inversion karyotype, and low values by species/region.

**Figure S12:**
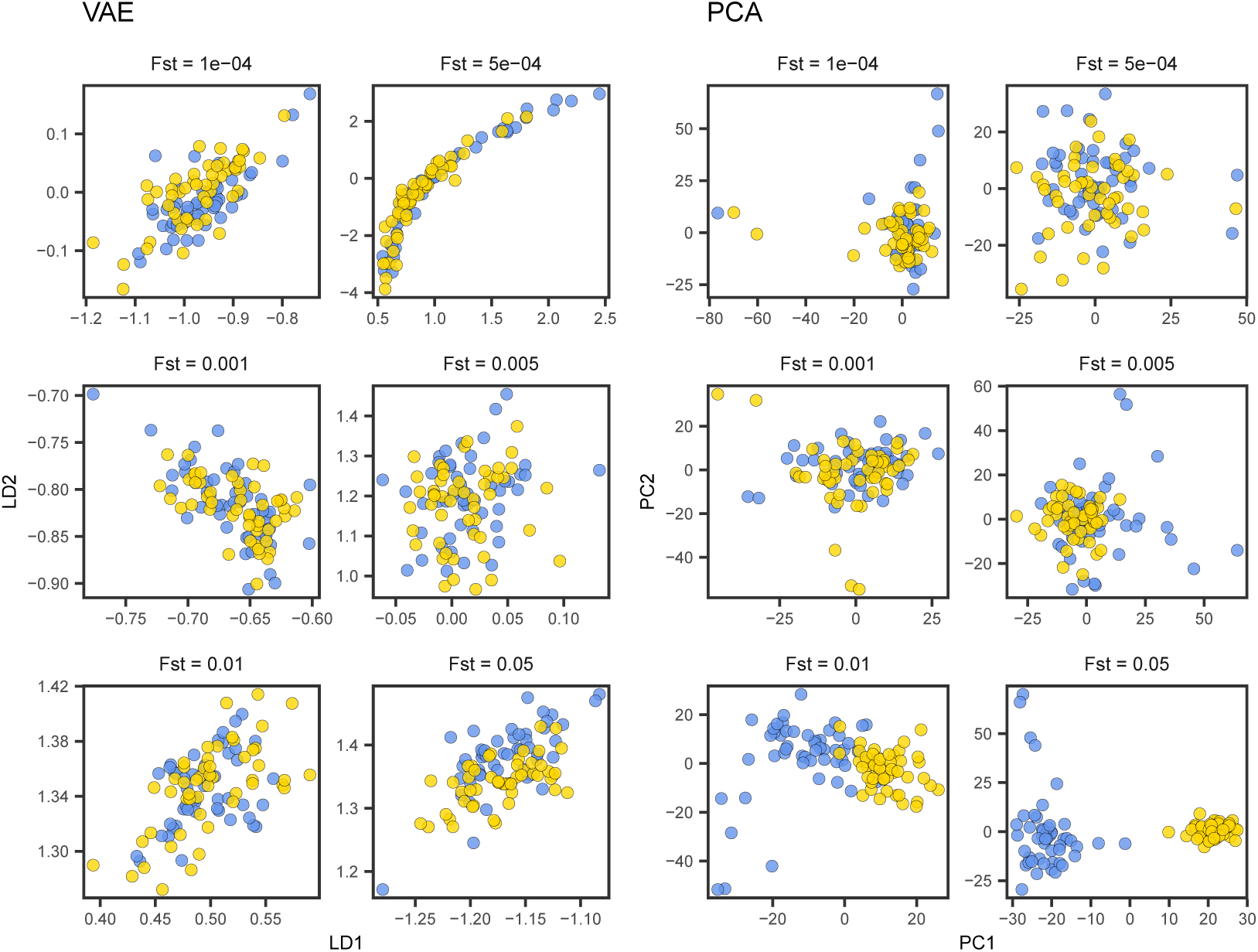
VAE latent spaces from the simulations shown in Figure 7, with `popvae` run at default settings.

**Figure S13:**
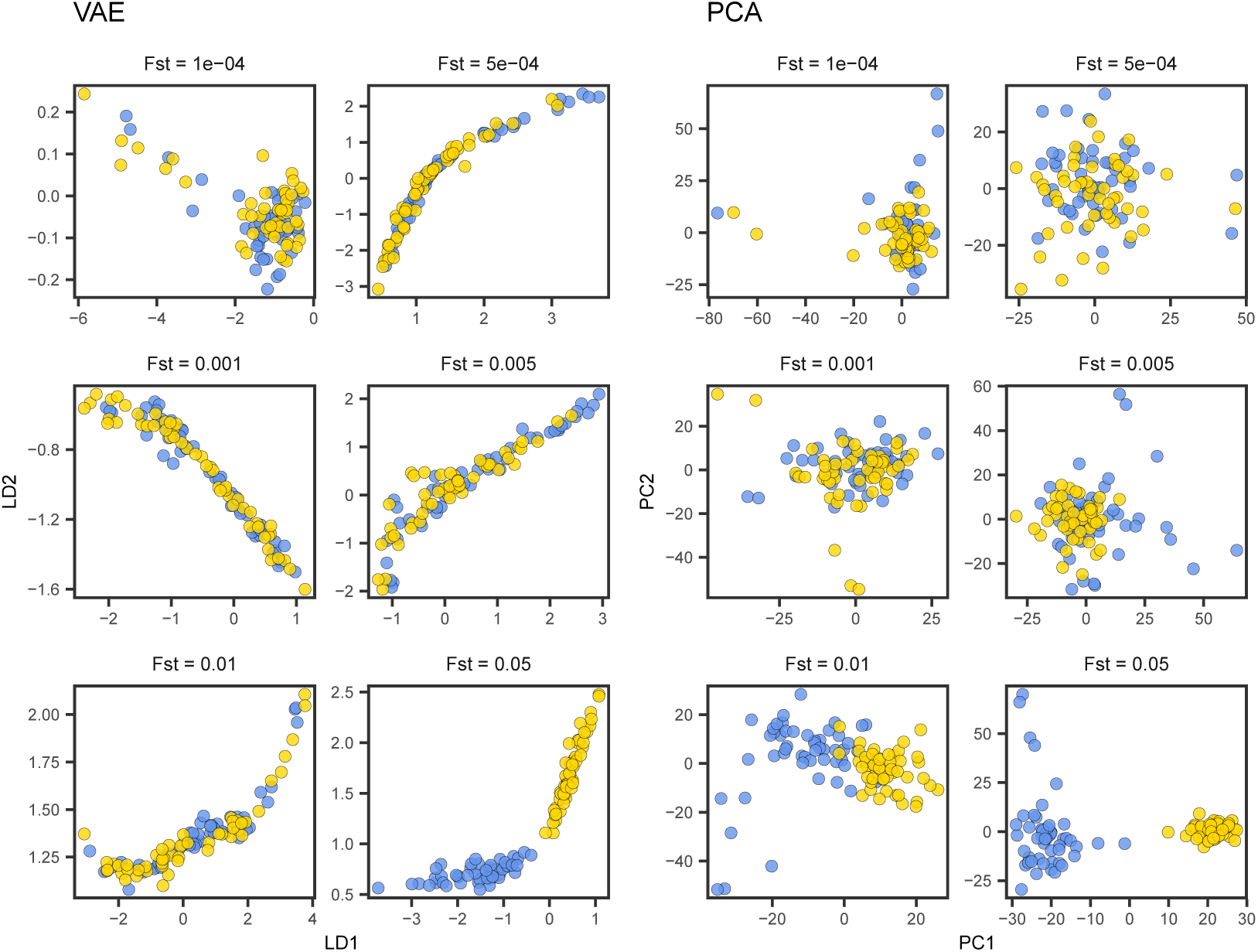
VAE latent spaces from the simulations shown in Figure 7, with `popvae` run with default network size (width 128, depth 6) and patience set to 500.

**Figure S14:**
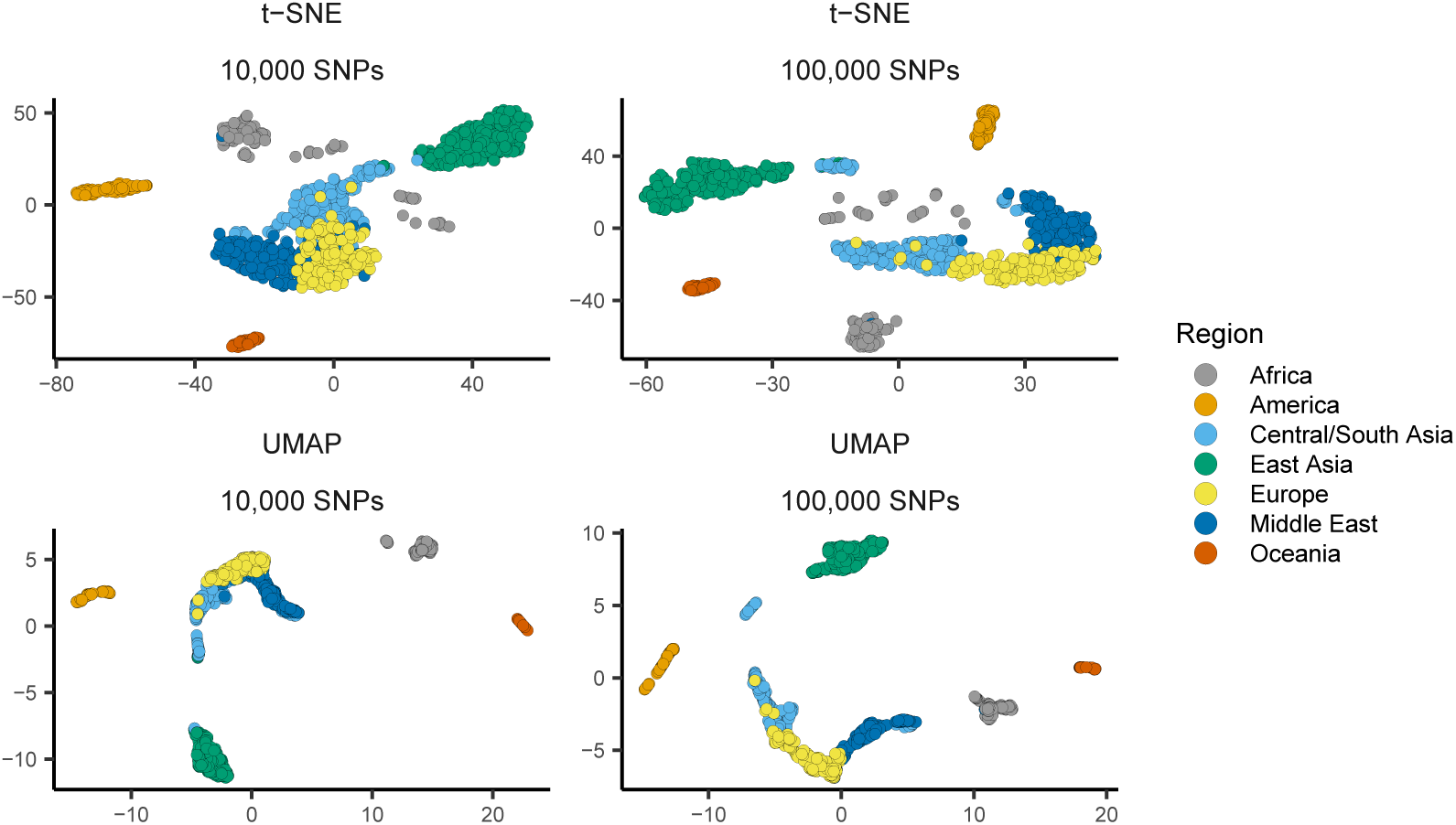
UMAP and t-SNE plots of HGDP samples using 100,000 or 10,000 SNPs. Both methods were run with default settings on the top 15 PC axes.

**Figure S15:**
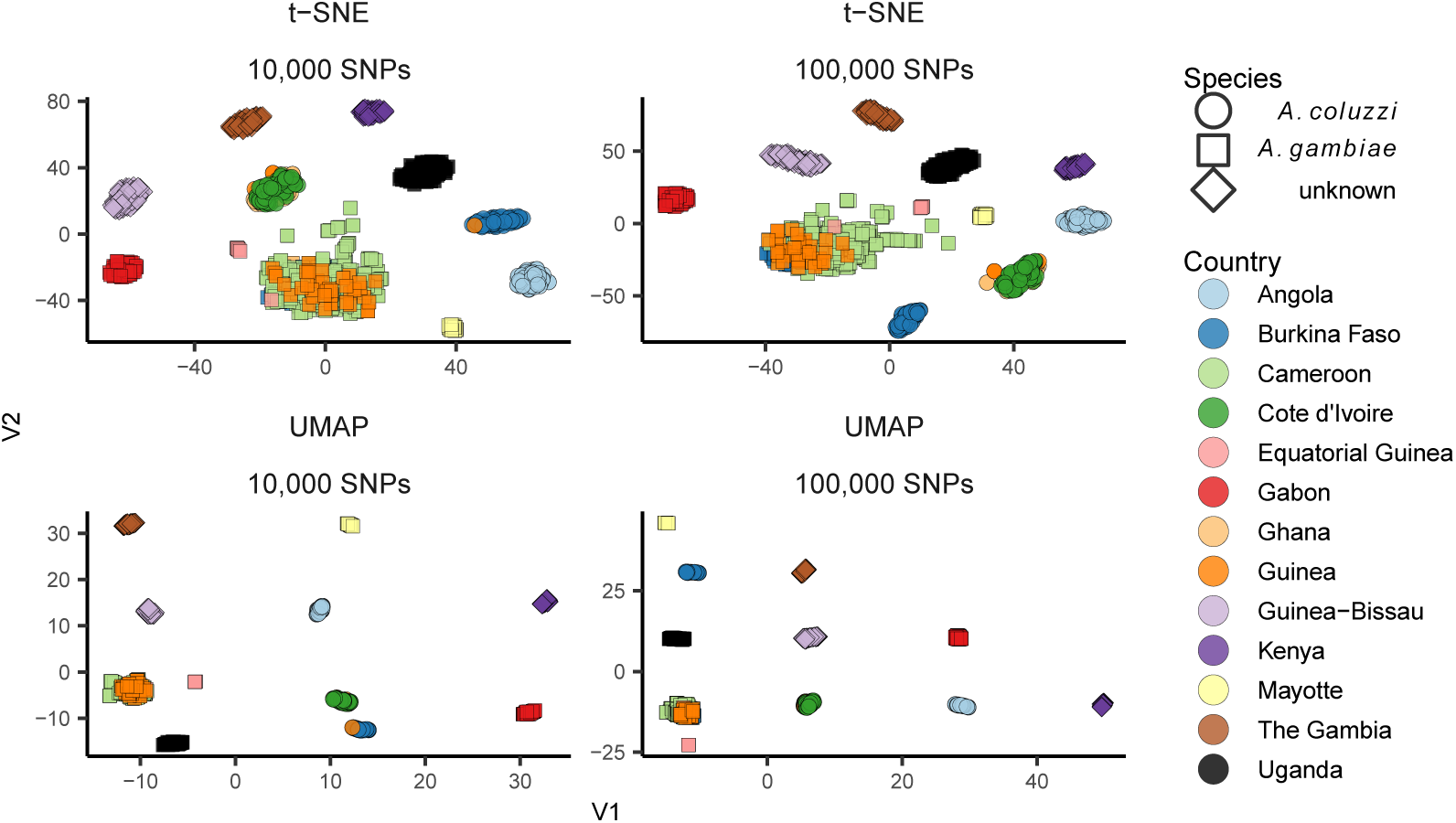
UMAP and t-SNE plots of AG1000G phase 2 samples using 100,000 or 10,000 SNPs at default settings. Both methods were run with default settings on the top 15 PC axes.

**Figure S16:**
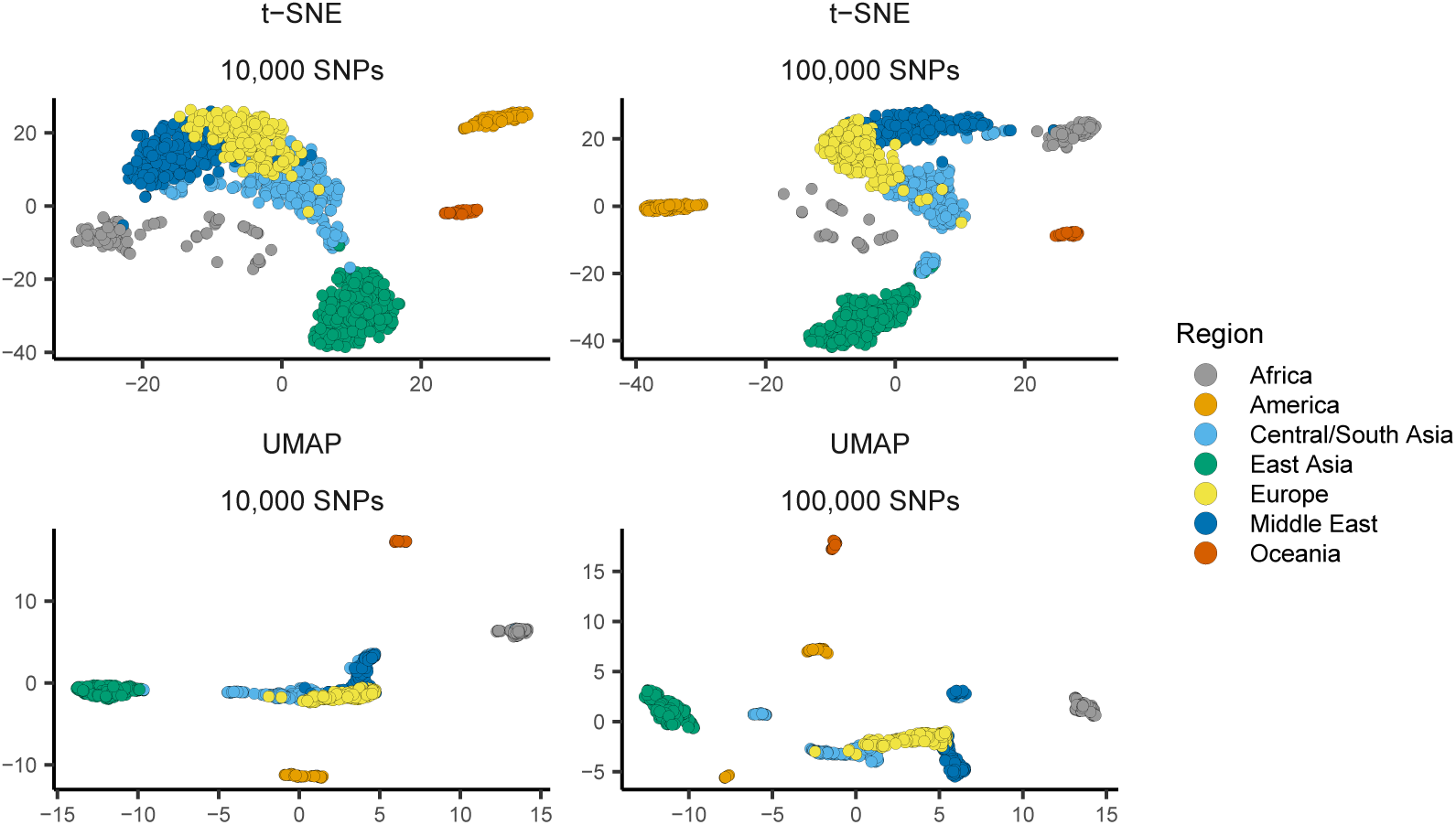
UMAP and t-SNE plots with parameters `n neighbors=30` and `perplexity=60`. These settings are double the default values and are intended to improve global relative to local structure.

**Figure S17:**
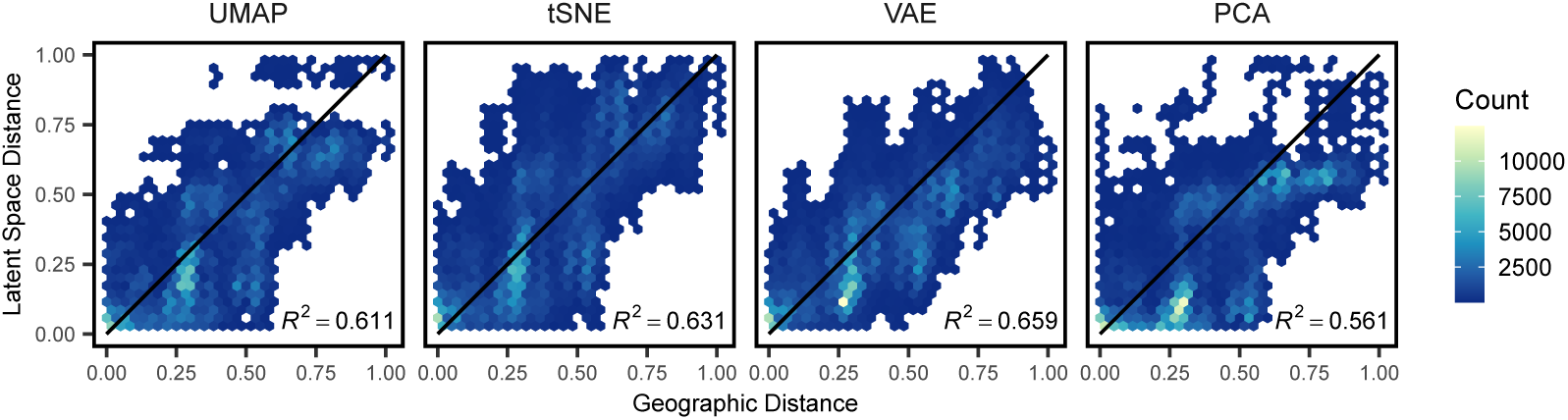
Comparison of relative pairwise distance for Eurasian HGDP samples, with UMAP parameter `n neighbors=30` and t-SNE parameter `perplexity=60`. These settings are double the default values and are intended to improve global relative to local structure.

**Figure S18:**
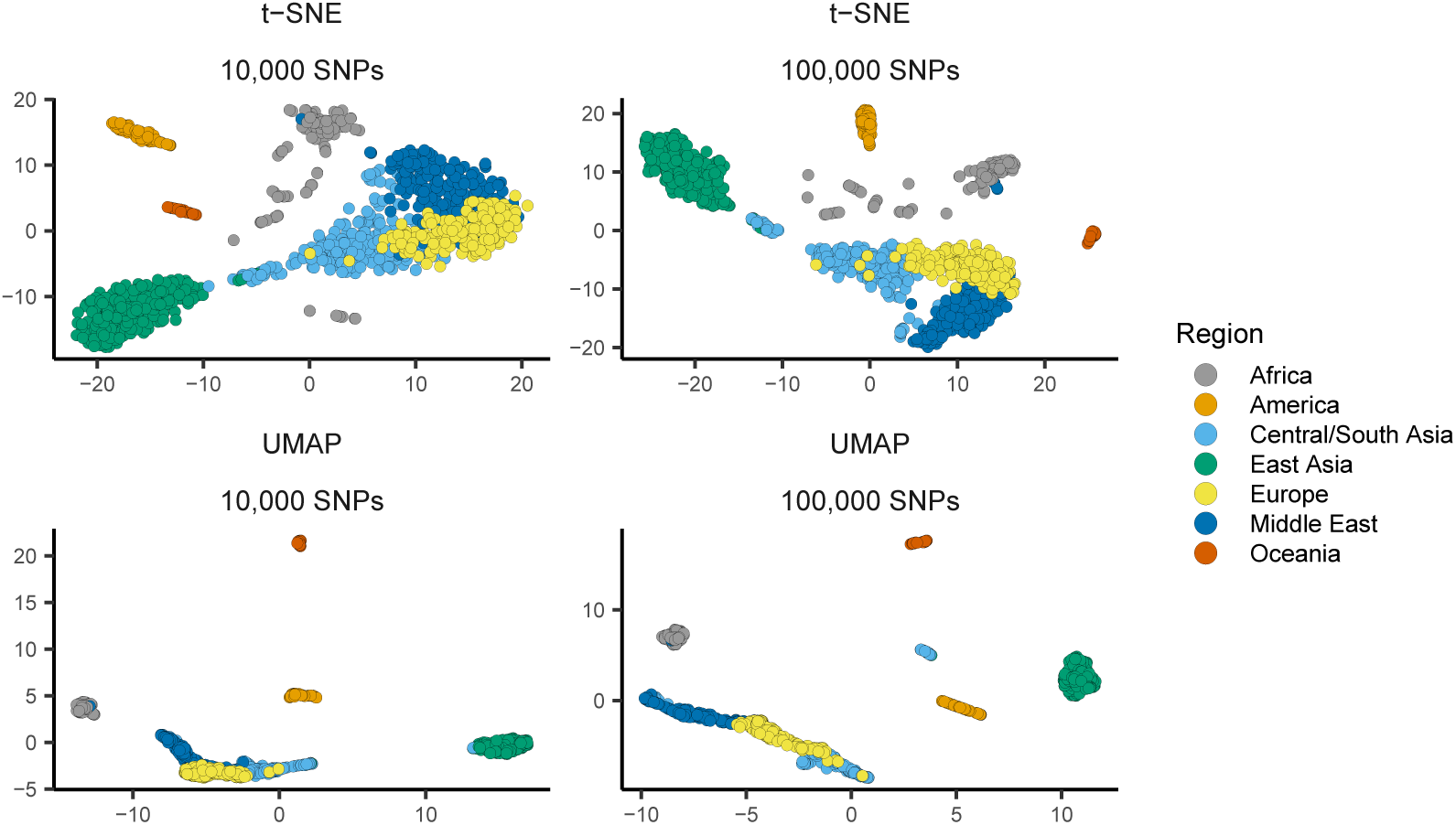
UMAP and t-SNE plots with parameters `n neighbors=30` and `perplexity=60`. These settings are double the default values and are intended to improve global relative to local structure.

**Figure S19:**
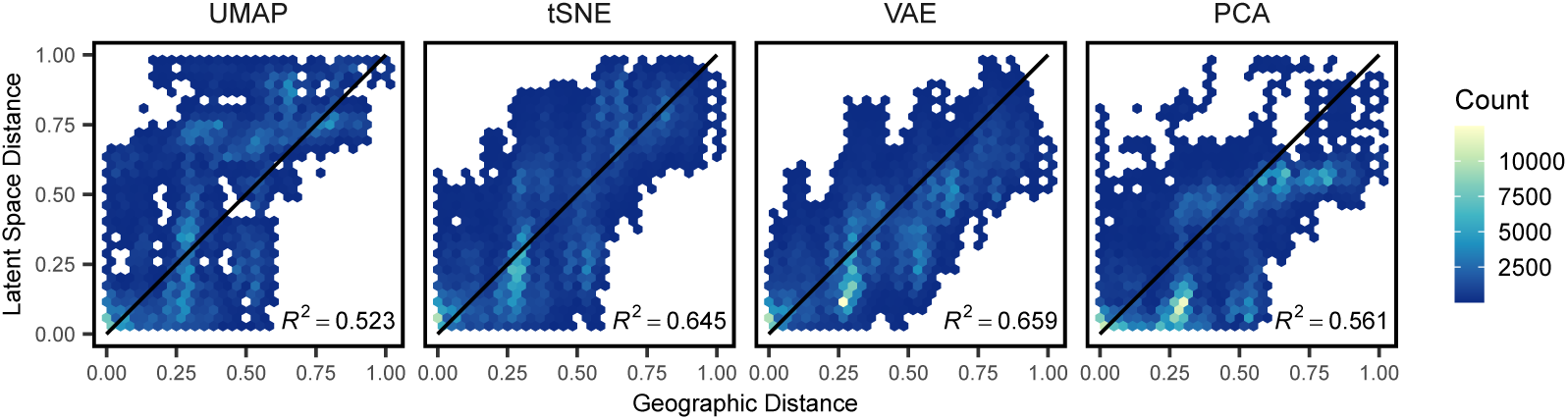
Comparison of relative pairwise distance for Eurasian HGDP samples, with UMAP parameter `n neighbors=45` and t-SNE parameter `perplexity=90`. These settings are triple the default values and are intended to improve global relative to local structure.

**Figure S20:**
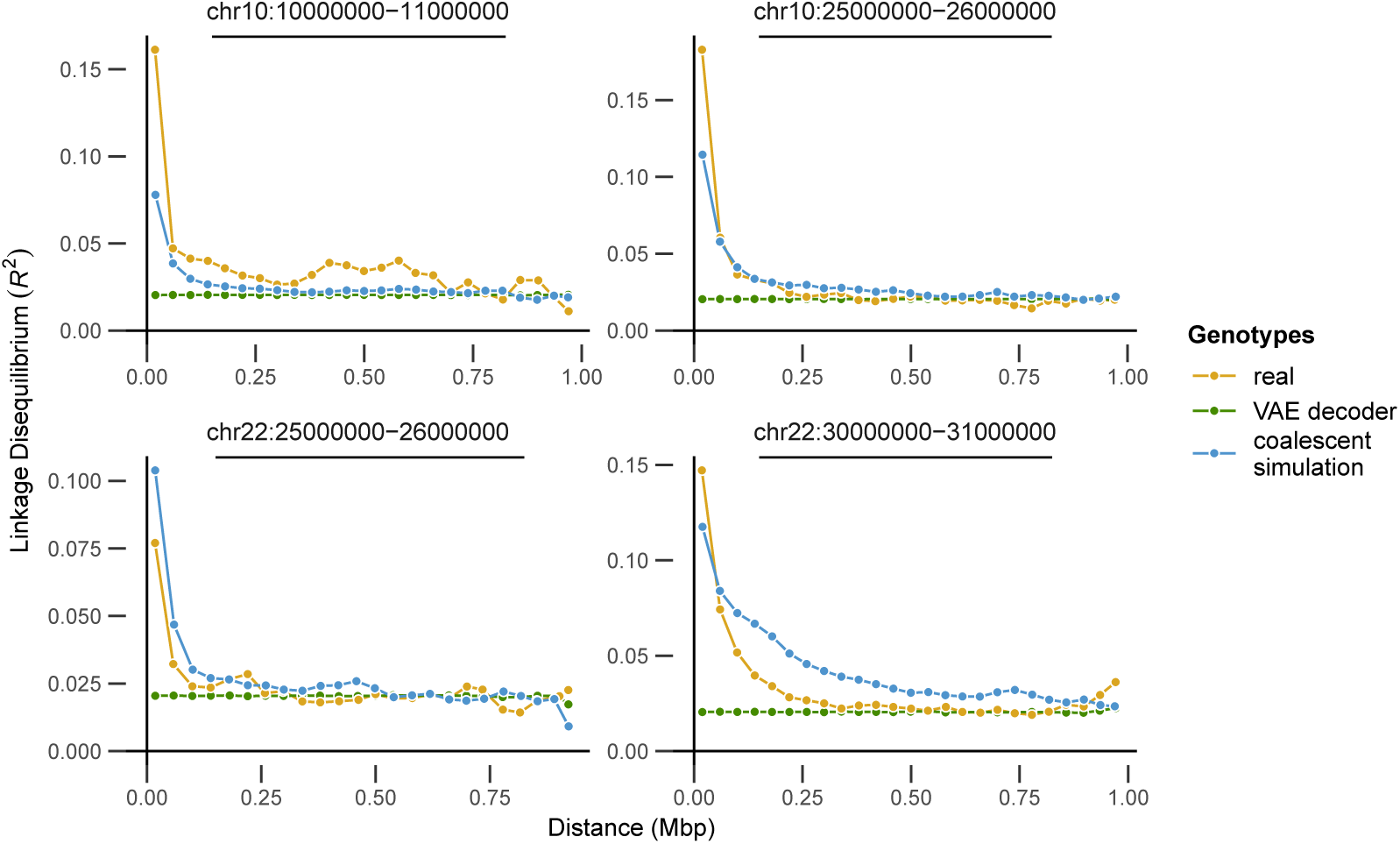
Comparing LD decay curves across real, simulated, and VAE decoder genotypes for four different regions of the genome. Points show the mean LD for all pairs of variants in each of 25 distance bins.

**Table S1:** Comparing validation loss across network sizes. Depth is the number of layers, width is the number of hidden units per layer, and loss is the mean validation loss across 5 random starting seeds for each network. Networks are ranked by loss for each dataset. SNPs were selected randomly from human chromosome 1 and *Anopheles* chromosome 3R.

| SNPs | HGDP |  |  | AG1000G |  |  |
| --- | --- | --- | --- | --- | --- | --- |
|  | Depth | Width | Loss | Depth | Width | Loss |
| **10,000** | 4 | 256 | 1394.231 | 3 | 256 | 750.677 |
|  | 6 | 128 | 1394.48 | 6 | 128 | 750.859 |
|  | 6 | 64 | 1394.504 | 4 | 128 | 751.0646 |
|  | 10 | 64 | 1394.663 | 4 | 256 | 751.1514 |
|  | 3 | 256 | 1394.976 | 6 | 64 | 751.7088 |
| **100,000** | 6 | 128 | 13955.76 | 4 | 256 | 7603.105 |
|  | 3 | 256 | 13968.39 | 6 | 128 | 7606.528 |
|  | 4 | 256 | 13971.75 | 3 | 256 | 7613.279 |
|  | 3 | 512 | 13980.04 | 6 | 256 | 7614.232 |
|  | 3 | 128 | 13992.27 | 4 | 128 | 7615.816 |
| **500,000** | 6 | 128 | 70087.32 | 10 | 128 | 37836.90 |
|  | 10 | 64 | 70191.43 | 6 | 128 | 37848.67 |
|  | 10 | 128 | 70203.22 | 6 | 64 | 37860.76 |
|  | 6 | 64 | 70221.66 | 10 | 64 | 37872.49 |
|  | 4 | 128 | 70357.73 | 4 | 128 | 37888.68 |

